# Efficient mixed model approach for large-scale genome-wide association studies of ordinal categorical phenotypes

**DOI:** 10.1101/2020.10.09.333146

**Authors:** Wenjian Bi, Wei Zhou, Rounak Dey, Bhramar Mukherjee, Joshua N Sampson, Seunggeun Lee

**Author notes:** Corresponding author: Wenjian Bi, Seunggeun Lee.

## Abstract

In genome-wide association studies (GWAS), ordinal categorical phenotypes are widely used to measure human behaviors, satisfaction, and preferences. However, due to the lack of analysis tools, methods designed for binary and quantitative traits have often been used inappropriately to analyze categorical phenotypes, which produces inflated type I error rates or is less powerful. To accurately model the dependence of an ordinal categorical phenotype on covariates, we propose an efficient mixed model association test, Proportional Odds Logistic Mixed Model (POLMM). POLMM is demonstrated to be computationally efficient to analyze large datasets with hundreds of thousands of genetic related samples, can control type I error rates at a stringent significance level regardless of the phenotypic distribution, and is more powerful than other alternative methods. We applied POLMM to 258 ordinal categorical phenotypes on array-genotypes and imputed samples from 408,961 individuals in UK Biobank. In total, we identified 5,885 genome-wide significant variants, of which 424 variants (7.2%) are rare variants with MAF < 0.01.

---

Large-scale biobanks with hundreds of thousands of genotyped and deeply phenotyped subjects are valuable resources to identify genetic components of complex phenotypes.^1,2^ In biobanks, ordinal categorical data is a common type of phenotype, which is often collected from surveys, questionnaires, and testing to measure human behaviors, satisfaction, and preferences.^3,4^ For example, a web questionnaire was used for 182,219 UK Biobank participants to collect 150 food and other health behavior related preferences, all of which are ordinal categorical phenotypes based on a 9-point hedonic scale of liking from 1 (extremely dislike) to 9 (extremely like).^5^ For ordinal categorical phenotypes, there is no underlying measurable scale and therefore it would be inappropriate to treat that phenotype as a quantitative trait and apply the linear regression methods.^6-8^ Another approach is to use an arbitrary cutoff to dichotomize the ordinal categorical phenotype into two categories, followed by using a logistic regression method.^3^ This approach suffers from information loss and thus is less powerful.

For binary and quantitative phenotype data analysis, mixed model approaches have been used to test genetic associations conditioning on the sample relatedness.^6,9^ Some state-of-art optimization strategies have been applied to reduce memory usage and computational cost, which makes these mixed model approaches practical to incorporate a dense genetic relationship matrix (GRM) in GWAS.^8,10^ Another resource-efficient approach, fastGWA, is to use sparse GRM to adjust for the sample relatedness.^11^ For binary phenotype analysis, unbalanced case-control ratio can result in inflated type I error rates and saddlepoint approximation (SPA) has been demonstrated to be more accurate for single-variant analysis^7,8^, region-based analysis^12,13^, and gene-environment interaction analysis^14^. Similarly, the sample size distribution in ordinal categorical data could also be highly unbalanced, that is, the sample size in one category could be dozens of times more than the that in other categories. For example, of the UK Biobank participants, more than 90% extremely dislike cigarette smoking and only 1% extremely like it. In ordinal categorical data analysis, the effect of the unbalanced sample size distribution on genetic association tests should also be carefully examined.

In this paper, we propose a scalable and accurate mixed model approach for ordinal categorical data analysis in large-scale GWAS. Our approach, Proportional Odds Logistic Mixed Model (POLMM), incorporates a random effect into the proportional odds logistic model to control for sample relatedness. POLMM uses penalized quasi-likelihood (PQL) and average information restricted maximum likelihood (AI-REML) algorithm^6^ to efficiently fit the mixed model, and then uses SPA to calibrate p values. We give two closely related versions, DensePOLMM and FastPOLMM. DensePOLMM incorporates a dense GRM using similar state-of-art strategies as in BOLT-LMM^10^ and SAIGE^8^, and FastPOLMM is a resource-efficient approach that uses sparse GRM in a similar manner as in fastGWA^11^.

We demonstrated that POLMM approaches can efficiently analyze large datasets with hundreds of thousands of genetic related samples, can control type I error rates, and is statistically powerful through extensive simulations as well as real data analysis. Meanwhile, BOLT-LMM, fastGWA, and SAIGE approaches cannot control type I error rates and are less powerful, especially when the phenotypic distribution is unbalanced. DensePOLMM requires comparable computation time and memory usage as SAIGE, and FastPOLMM is more resource-efficient to fit a null mixed model. For example, FastPOLMM requires less than 0.1 hour and 4.2 GB memory to fit a null mixed model with around 400,000 subjects. In most scenarios, DesnePOLMM and FastPOLMM performed similarly. Only when the number of categories is large (e.g. 10) and polygenic effect size is large (e.g. liability heritability = 75.24%), DensePOLMM is slightly more powerful than FastPOLMM by no more than 4.67% and 7.51% when testing common (minor allele frequency, MAF = 0.3) and low-frequency variants (MAF = 0.01), respectively. We applied the FastPOLMM approach to analyze 258 ordinal categorical phenotypes in the UK Biobank data, which includes 408,961 samples from white British participants with European ancestry, and successfully identified 5,885 genome-wide significant variants with clumping, of which 424 variants (7.2%) are rare variants with MAF < 0.01. All analysis results have been publicly available through a web-based visual server^2^, which provides intuitive visualizations at three levels of granularity: genome-wide summaries at the trait level, and regional (LocusZoom)^15^ and phenome-wide summaries at the variant level.

## Results

### Overview of the methods

We let *n* denote the sample size and let *J* denote the number of category levels. For subject *i* ≤ *n*, we let *y*_*i*_ *=* 1,2,…, *J* denote its ordinal categorical phenotype. We consider the following proportional odds logistic mixed model (POLMM)

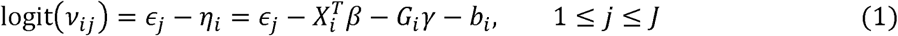

where *v*_*i,j*_ *=* Pr(*y*_*i*_ ≤ *j*|*X*_*i*_, *G*_*i*_, *b*_*i*_) is the cumulative probability of the phenotype *y*_*i*_ ≤ *j* conditional on a *p-*dimensional vector of covariates *X*_*i*_ and a hard called or imputed genotype *G*_*i*_. The cutpoints *ϵ: ϵ*_1_ *<* … *< ϵ* _*j*_ *=* ∞ were used to categorize the data, and coefficients *β* and *γ* are fixed effect sizes of the covariates and genotype. To adjust for sample relatedness, we incorporate an *n*-dimensional random effect vector *b = (b*_1_, …, *b*_*n*_)^*T*^ following a multivariate normal distribution *N*(0, *τV*) where *τ* is a variance component parameter and *V* is an *n* × *n* dimensional GRM. The model (1) is a natural extension of a logistic mixed model as in SAIGE and GMMAT.^6,8,13^ If *J* = 2, the phenotype is binary and the model (1) is a logistic mixed model.

We present two closely related versions of POLMM methods to test null model *γ* = 0: DensePOLMM and FastPOLMM. The methods contain two main steps: (1) fitting the null model to estimate the variance component 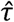 and other parameters 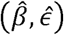; (2) testing for the association between the ordinal categorical phenotype and genetic variants. In step 1, we propose an efficient algorithm (Supplementary Note) to iteratively estimate parameters using PQL and AI-REML algorithm^6^. DensePOLMM and FastPOLMM use dense and sparse GRM to adjust for sample relatedness, respectively. DensePOLMM stores raw genotypes in a binary vector, calculates elements of the dense GRM when needed, and uses PCG approach to solve linear systems.^8^ Using these strategies, DensePOLMM is of the same computation complexity as SAIGE^8^, and requires memory usage *m*_1_*n/*4, where *m*_1_ is the number of markers used to construct GRM. On the other hand, FastPOLMM uses a sparse GRM in which all of the small off-diagonal elements (for example, those < 0.05) are set to 0. GCTA software^16^ provides an efficient tool to calculate GRM for a large-scale dataset. The sparse GRM only needs to be calculated once for one cohort study or biobank.

After fitting the null mixed model, we first use a subset of randomly selected genetic variants to calculate the ratio of the variances of the score statistics with and without incorporating the variance component. The ratio has been shown approximately constant for all genetic variants with minor allele counts (MAC) ≥ 20.^8,10^ Then, we use the variance ratio to calibrate the score statistic variance (Supplementary Note). To control type I error rates under an unbalanced phenotypic distribution, we use a hybrid strategy of normal approximation and SPA to calculate p values.^7,8,14,17^ If the absolute value of the standardized score statistic is close to the mean of 0 (e.g. < 2), POLMM methods use a regular normal approximation to calculate p value. Otherwise, POLMM methods use SPA to calculate p values. The hybrid strategy can give more accurate p values while remaining high computationally efficient. For each variant, the normal approximation takes *O(n)* computations and SPA additionally takes *O*(*n*(*J* − 1)) computations. Using the fact that many elements of the genotype vector *G = (G*_1_, …, *G*_*n*_)^*T*^ are zeros (i.e., homozygous major genotypes), we use a partial normal approximation^7^ to speed up the computation of SPA to *O*(*n*_1_(*J* − 1)) where *n*_1_ is the number of non-zero elements in the genotype vector *G*.

Due to these features, POLMM methods are the only available mixed model methods to associate ordinal categorical phenotypes with genetic variants while remaining computationally practical for large datasets and accounting for sample relatedness and unbalanced phenotypic distribution.

### Runtime and resource requirements

To evaluate the computational efficiency and memory usage of DensePOLMM and FastPOLMM methods, we randomly sampled subjects from 397,798 white British UK Biobank participants to analyze an ordinal categorical phenotype, able to confide, which consists of 6 levels (Figure S1). We used 340,447 markers to construct GRM and incorporated 6 covariates of sex, birth year, and top 4 principal components to fit the null mixed model.

We compared 5 methods including fastGWA, BOLT-LMM, SAIGE, DensePOLMM, and FastPOLMM. Besides the raw phenotype with 6 levels, we combined some levels to make a new phenotype with 3 levels to comprehensively evaluate POLMM methods. For fastGWA and BOLT-LMM, we treated the ordinal categorical phenotype as a quantitative trait from 1 to 6. For SAIGE, we dichotomized the phenotype to a binary phenotype. For fastGWA and FastPOLMM, we set the cutoff of the sparse GRM at 0.05. Details about the computing environment for evaluation can be seen in Methods.

The computation time and memory usage of all 5 methods are presented in Figure S2 and Table S1. In step 1, to fit a null mixed model, fastGWA and FastPOLMM were much faster and required much less memory than the three methods using dense GRM. BOLT-LMM, SAIGE, and DensePOLMM required comparable computation time and memory usage since they used the same optimized strategies to incorporate the dense GRM. SAIGE and DensePOLMM were slower than BOLT-LMM since they use Hutchinson’s randomized trace estimator when estimating the variance component, which requires a large amount of computation time. DensePOLMM required more time than SAIGE when sample size was greater than 100,000. This is mainly because DensePOLMM used a block diagonal matrix as the preconditioner matrix for PCG, which took more iterations to converge than that in SAIGE given the same tolerance criterion. Interestingly, DensePOLMM was faster than SAIGE when the sample size was smaller than 40,000. This might be because we optimized C++ codes to read in genotypes for GRM construction. For POLMM methods, more computational time and slightly more memory usage were required when analyzing a phenotype with more levels. For example, to fit a null mixed model with 397,798 subjects, if the number of levels is 3, DensePOLMM and FastPOLMM took 49.9 and 0.03 hours, respectively; and if the number of levels is 6, DensePOLMM and FastPOLMM took 64.2 and 0.09 hours, respectively.

In step 2, we first recorded the computation time to analyze 340,447 markers and then projected them to a genome-wide analysis with 30 million markers. The genotype data was stored in BGEN format since all methods in comparison support the BGEN format and UK Biobank also use it for release of imputed data.^18^ BOLT-LMM and fastGWA were faster than POLMM and SAIGE methods, which is expected as logistic regression is more complicated than linear regression. POLMM is slightly faster than SAIGE. As the number of levels increased from 3 to 6, the computation time of POLMM methods slightly increased. Suppose that we use 24 CPU cores for parallel computation, POLMM methods require around 14.2 hours for a genome-wide analysis including around 30 million markers.

### False positive rate and statistical power

We carried out extensive simulations to investigate type I error rates and powers of POLMM approaches. We simulated 10,000 subjects in 1,000 families based on the pedigree shown in Figure S3, in which each family included 10 subjects. To construct GRM for mixed model methods, we simulated 100,000 single nucleotide polymorphisms (SNPs) with MAFs ranging from 0.05 to 0.5. The estimated kinship coefficients are shown in Figure S4. We simulated phenotypes with multiple sample size distributions (Figure S5). In addition to DensePOLMM and FastPOLMM that use a hybrid of normal distribution approximation and SPA, we also evaluated DensePOLMM-NoSPA and FastPOLMM-NoSPA, both of which use normal distribution approximation to test all variants. We also evaluated some alternative methods including SAIGE, fastGWA, and BOLT-LMM. For SAIGE, we dichotomized the categorical phenotypes (Figure S5). For fastGWA and BOLT-LMM, we treated the categorical phenotype as a quantitative trait.

We first simulated categorical phenotypes under the null model to evaluate type I error rates. In each scenario, a total of 10^9^ tests were performed (Methods). The simulation results showed that DensePOLMM and FastPOLMM methods can control type I error rates at a significance level of 5 × 10^−8^ (Figures 1 and S6). Meanwhile, type I error rates of other methods were inflated when testing low-frequency and rare variants (MAF ≤ 0.01) and the phenotypic distribution was unbalanced. For example, when variance component was *τ* = 1 and the sample size proportion in 4 levels was 100:1:1:1, to test low-frequency variants with a MAF of 0.01, the type I error rates of POLMM methods and the other methods were less than 3.8 × 10^−8^ and greater than 3.89 × 10^−6^, respectively. The result suggested that POLMM approaches can accurately account for ordinal categorical responses and using SPA is more accurate than using normal distribution. If we dichotomize the categorical phenotype, the POLMM is a logistic mixed model and it is expected that SAIGE can control type I error rates.^8^ Hence, we did not evaluate the empirical type I error rates of SAIGE.

**Figure 1.**
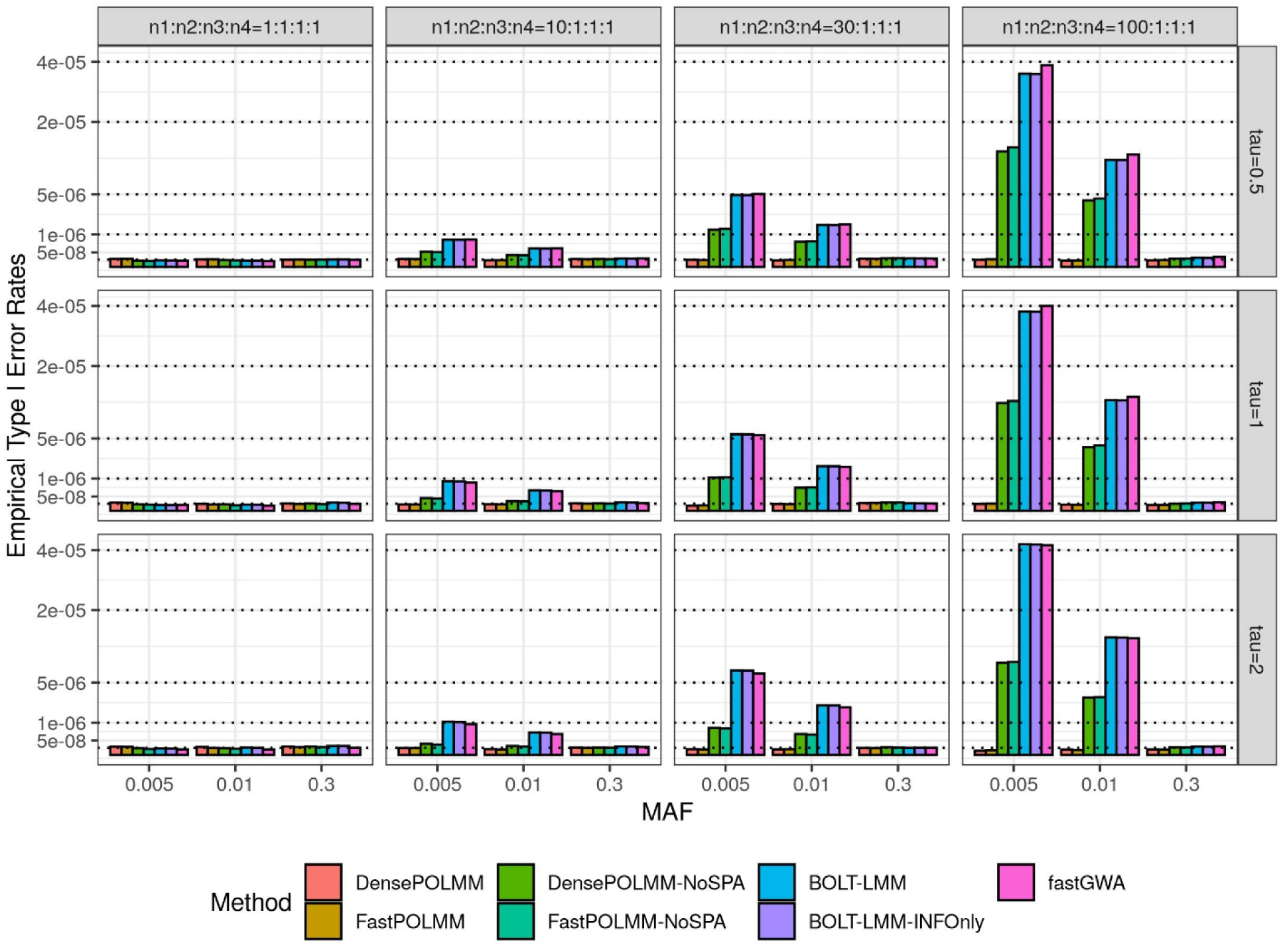
Empirical type I error rates of POLMM, BOLT-LMM, and fastGWA methods at a significance level 5e-8. We simulated 1,000 families with a total sample size *n =* 10,000 and an ordinal categorical phenotype including four levels with sample sizes *n*_1_, *n*_2_, *n*_3_, and *n*_4_. From left to right, the plots consider four scenarios: balanced (*n*_1_: *n*_2_: *n*_3_:*n*_4_ *=* 1:1:1:1), moderately unbalanced (*n*_1_: *n*_2_: *n*_3_:*n*_4_ *=* 10:1:1: 1), unbalanced (*n*_1_:*n*_2_:*n*_3_:*n*_4_ = 30:1:1:1), and extremely unbalanced (*n*_1_. *n*_2_: *n*_3_: *n*_4_ *=* 100:1:1:1). From top to bottom, the plots consider three variance components *τ* = 0.5, 1, and 2. We simulated common, low-frequency, and rare variants with MAFs of 0.3, 0.01 and 0.005, respectively.

Next, we compared the empirical powers of POLMM methods, SAIGE, fastGWA, and BOLT-LMM at a significance level *α =* 5 × 10^−8^ (Figures 2 and S7). Since fastGWA and BOLT-LMM cannot control type I error rates when the phenotypic distribution is unbalanced, we used empirical significance levels to evaluate powers. In all simulation scenarios, POLMM methods were the most powerful. When the phenotypic distribution is balanced, fastGWA and BOLT-LMM were similarly powerful as POLMM methods. However, when the phenotypic distribution is unbalanced, fastGWA and BOLT-LMM methods were less powerful than POLMM methods, especially when testing low-frequency variants with MAF = 0.01. Since the dichotomizing process would result in information loss, SAIGE was also less powerful than POLMM methods. Figure S7 showed that different dichotomizing processes could result in significantly different powers for SAIGE.

**Figure 2.**
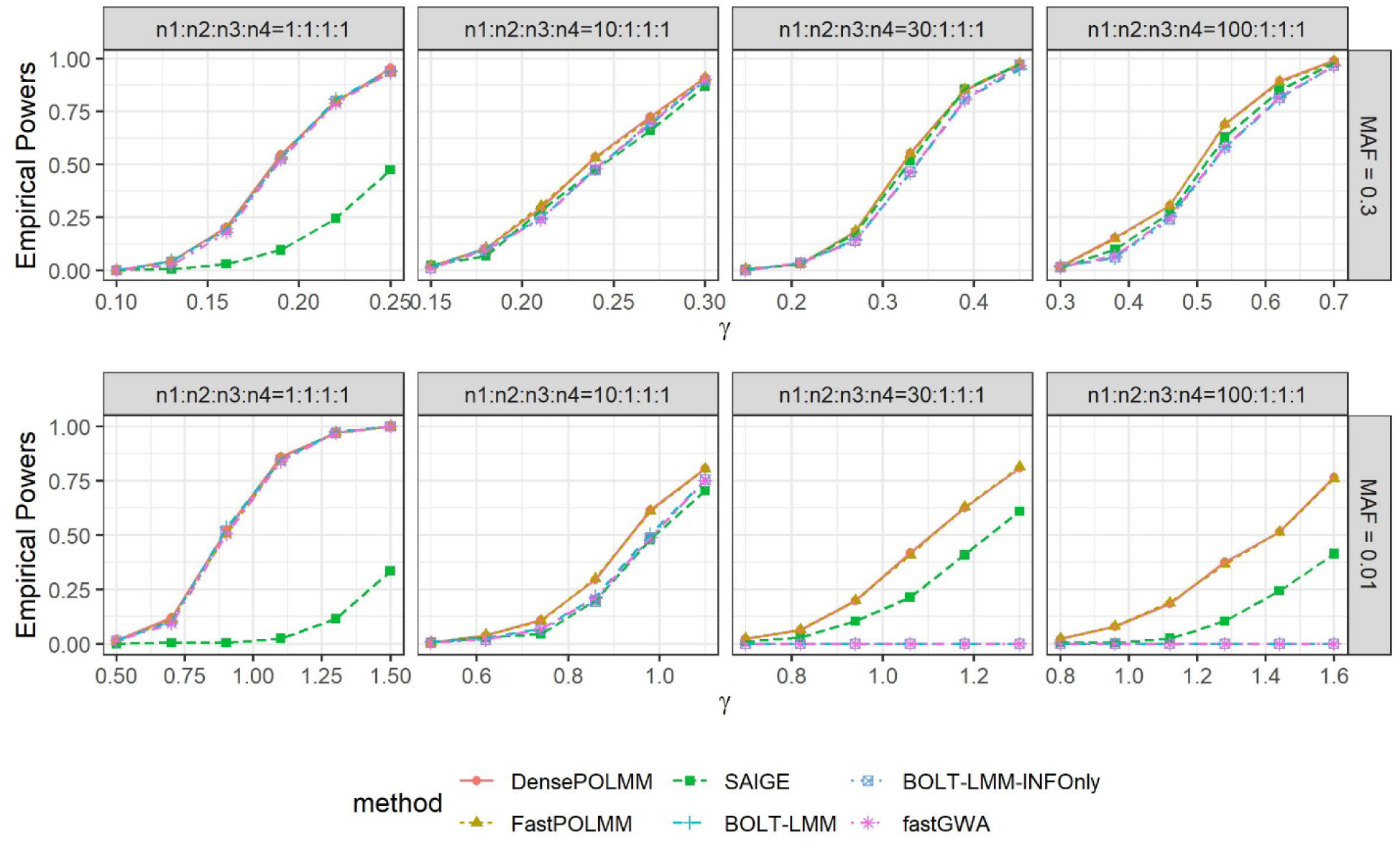
Empirical powers of POLMM, SAIGE, BOLT-LMM, and fastGWA methods at significance level 5e-8. We simulated 1,000 families with a total sample size *n =* 10,000 and an ordinal categorical phenotype including four levels with sample sizes *n*_1_, *n*_2_, *n*_3_, and *n*_4_. From left to right, the plots consider four scenarios: balanced (*n*_1_. *n*_2_. *n*_3_:*n*_4_ *=* 1:1:1:1), moderately unbalanced (*n*_1_. *n*_2_. *n*_3_:*n*_4_ *=* 10:1:1: 1), unbalanced (*n*_1_: *n*_2_:*n*_3_: *n*_4_ *=* 30:1:1:1), and extremely unbalanced (*n*_1_. *n*_2_: *n*_3_:*n*_4_ *=* 100:1:1:1). From top to bottom, the plots consider two MAFs of 0.3 and 0.01 to simulate common and low-frequency variants. We let variance component *τ*= 1. For SAIGE, we dichotomize phenotype as 0 or 1 depending on the subject is in level 1 or not. For BOLT-LMM, the empirical powers were calculated based on the empirical significance levels since it cannot control type I error rates for low-frequency variants.

### Comparison between DensePOLMM and FastPOLMM methods

For quantitative trait analysis, Jiang et al. have demonstrated that using sparse GRM can reduce computational time and memory usage, while still being reliable to control type I error rates.^11^ However, using spare GRM can be less powerful than using dense GRM since sparse GRM cannot incorporate polygenic effects. In this section, we designed more simulation scenarios to compare DensePOLMM and FastPOLMM (Methods).

Figures S8-S11 present the variance component estimation 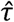 and the empirical powers of POLMM methods. The estimation 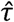 of DensePOLMM and FastPOLMM were slightly different, both of which deviated from true *τ*, especially when the true *τ* was large. The biased estimation has been widely discussed in other studies using pseudo quasi likelihood (PQL).^8^ Interestingly, the estimation 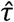 increased and tended to the true *τ* as the number of levels increased from 3 to 10. This might be because more levels give more information, which results in more accurate estimation of the variance component *τ*. In most scenarios, the empirical powers of DensePOLMM and FastPOLMM were similar with the largest difference less than 2.5%. Only when SNPs used to construct GRM were significantly associated with the phenotype (e.g. liability heritability = 75.24%, see Methods) and the number of levels is large (e.g. 10), DensePOLMM is more powerful than FastPOLMM by no more than 4.67% and 7.51%, when testing SNPs with MAF = 0.3 and 0.01, respectively. This may be because that only when the number of levels is large, accounting for the polygenic effects through dense GRM can substantially improve the power. Note that in this simulation, SNPs for dense GRM were simulated independently from the SNPs to test, to prevent proximal contamination.

Compared to DensePOLMM, FastPOLMM can give a substantial improvement in terms of computation time and memory usage, while only suffering a limited loss of powers in restricted simulation scenarios. Hence, we recommended using FastPOLMM, especially when analyzing a large-scale dataset with sample size greater than 200,000.

### Application to UK Biobank Data

We used FastPOLMM to conduct genome-wide analyses of 30 million SNPs with minor allele counts ≥ 20 and imputation R^2^ greater than 0.3 in the UK Biobank data of 408,961 samples from white British participants. We incorporated birth year, sex (if applicable), and top 4 principal components as covariates, and used 340,447 high-quality SNPs to calculate sparse GRM in which all of off-diagonal elements less than 0.05 were set to 0.^8,16^ We analyzed 258 ordinal categorical phenotypes, most of which measured dietary, lifestyle and environment, and psychosocial factors (Table S2). All analysis results have been publicly available through a visual server (http://polmm.leelabsg.org/). The web interface provides intuitive visualizations at three levels of granularity: genome-wide summaries at the trait level, and regional (LocusZoom)^15^ and phenome-wide summaries at the variant level.^2^

To compare BOLT-LMM and FastPOLMM in ordinal categorical data analysis, we selected four food preferences with different sample size distribution as phenotypes (Figure S12). The preferences were encoded from 1 (extremely dislike) to 9 (extremely like). For BOLT-LMM, we treated the phenotypes as quantitative traits and incorporated the same set of covariates and GRM as in FastPOLMM. Figures 3 and S13 present the Manhattan and QQ plots of the analysis results. When the phenotypic distribution is balanced, BOLT-LMM performed similarly as FastPOLMM. However, in other cases, BOLT-LMM could result in an inflation of type I error rates, especially when testing low-frequency and rare variants with MAF < 0.01. FastPOLMM-NoSPA was better than BOLT-LMM but still cannot control type I error rates at a genome-wide significance level, which suggests that the proportional odds logistic model and SPA both contribute to more accurate association tests. All the real data analysis results were consistent to the simulation results, which indicate that using linear models is not a good solution in ordinal categorical data analysis, especially when testing low-frequency variants.

**Figure 3.**
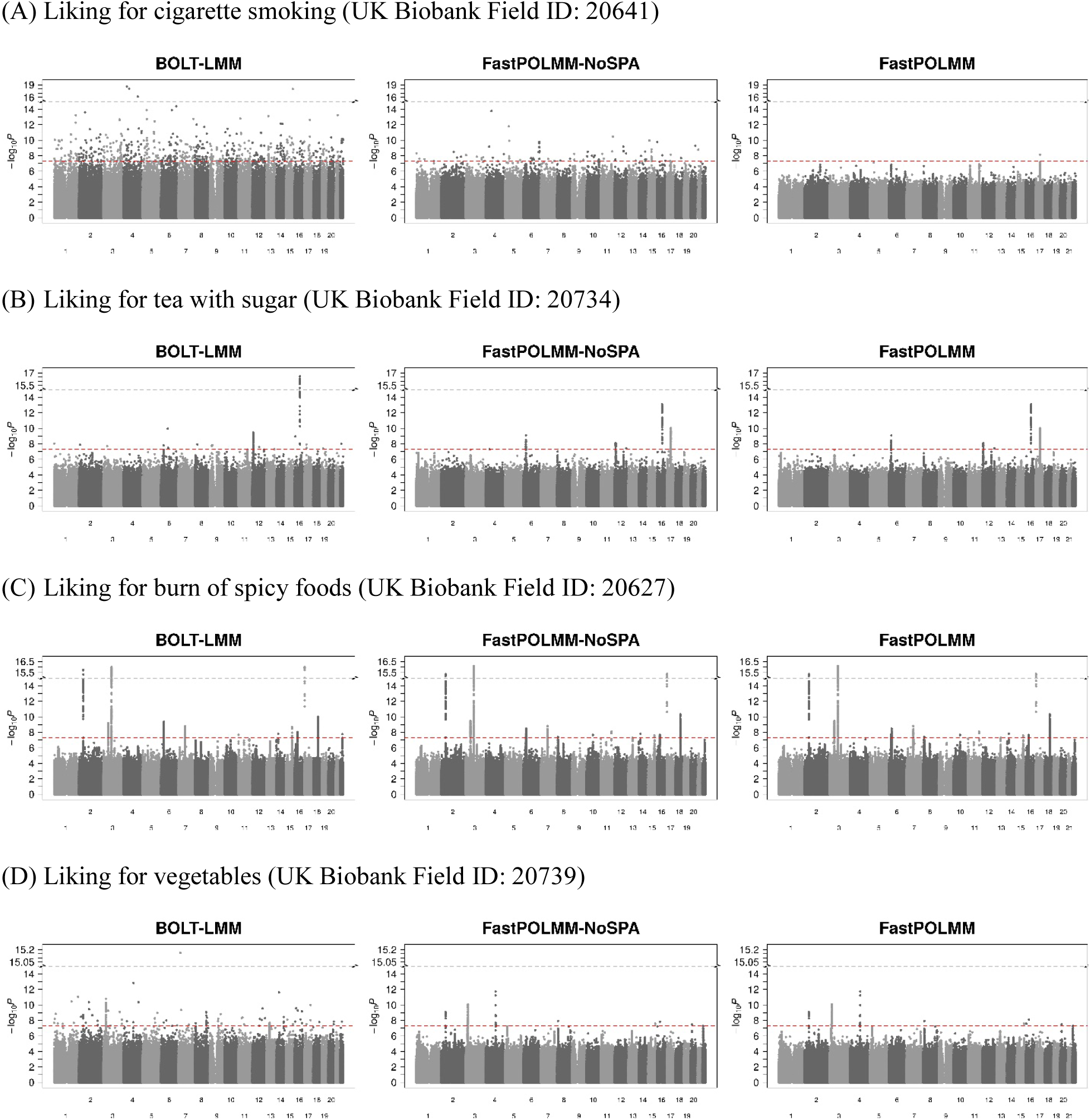
Manhattan plots for UK Biobank analysis

We used PLINK^19^ to conduct clumping analysis (p value threshold of 5 × 10^−8^, window size of 5 Mb, and linkage disequilibrium threshold *r*^2^ of 0.1). For these 258 phenotypes, we identified 5,885 clumped genome-wide significant variants, of which 424 variants (7.2%) are low-frequency variants with MAF < 0.01. We used ANNOVAR^20^ to functionally annotate these genome-wide significant variants. Total 275 clumped variants are in exon region, of which 207 (75.3%, binomial test p value: 1.04E-12) variants are nonsynonymous variants. Based on the Polyphen2 HDIV score^21^, 63 nonsynonymous variants (30.4%, binomial test p value: 0.506) are probably damaging with the score ≥ 0.957 and 33 nonsynonymous variants (15.9%, binomial test p value: 1) are possibly damaging with the score ≥ 0.453. Table S3 gives a summary of the annotation of more than 24 million SNPs, which were used to calculate the proportion of nonsynonymous variants, probably damaging variants, and possibly damaging variants.

We highlight some nonsynonymous low-frequency SNPs with significant associations. For phenotype of “morning/evening person” (UK Biobank Field ID: 1180), we identified an association of nonsynonymous SNP rs139315125 (MAF: 0.47%, p value: 5.3E-21, Gene: *PER3*, Polyphen2 HDIV score: 0.998, see Figure S14 for more details). Subjects who tend to sleep and wake up early have a higher frequency of minor allele G. Gene *PER3* is a core component of the circadian clock and the association between this SNP and sleep-wake patterns has been reported in previous studies.^22^ For phenotype of “Use of sun/uv protection” (UK Biobank Field ID: 2267), we identified a nonsynonymous SNP rs121918166 (MAF: 0.9%, p value: 5.2E-31, Gene: *OCA2*, Polyphen2 HDIV score: 1, see Figure S15 for more details). Subjects who use sun/uv proection more frequently have a higher frequency of minor allele T. Gene *OCA2* is involved in mammalian pigmentation and this SNP has been previously associated with human eye color and melanoma.^23-25^ Other interesting association include phenotype of “Comparative height size at age 10” (UK Biobank Field ID: 1697) and rs78727187 (MAF: 0.6%, p value: 5.1E-19, Gene: *FBN2*, Polyphen2 HDIV score: 0.818), rs117116488 (MAF: 0.99%, p value: 1.4E-18, Gene: *ACAN*, Polyphen2 HDIV score: 0.993), and rs112892337 (MAF: 0.4%, p value: 3.0E-15, Gene: *ZFAT*, Polyphen2 HDIV score: 1); phenotype of “Relative age of first facial hair” (UK Biobank Field ID: 2375) and rs138800983 (MAF: 0.3%, p value: 8.4E-10, Gene: *KRT75*, Polyphen2 HDIV score: 0.969).

## Discussion

In this study, we developed a scalable and accurate genetic association analysis tool, POLMM, for ordinal categorical data analysis in a large-scale dataset with hundreds of thousands of samples. The tool is an extension of proportional odds logistic model, which can accurately account for the dependence of an ordinal categorical phenotype on covariates. Two closely related methods, DensePOLMM and FastPOLMM, were proposed to use dense and sparse GRM to adjust for the sample relatedness, respectively. DensePOLMM uses similar optimized strategies as in SAIGE and BOLT-LMM, which makes it scalable to incorporate a dense GRM into the mixed model. However, as the sample size increases, DensePOLMM is still computationally expensive. On the other hand, FastPOLMM is more computationally efficient. Extensive simulations demonstrate that FastPOLMM is as reliable as DensePOLMM and only suffers a small amount of power loss in limited simulation scenarios. Hence, if the sample size is greater than 500,000 and hundreds of GWAS are required for a phenome-wide analysis, we recommend using FastPOLMM.

We compared our method POLMM with two commonly used strategies: 1) dichotomize the categorical phenotype and then use SAIGE^8^, and 2) treat the categorical phenotype as a quantitative trait and then use BOLT-LMM^10^ and fastGWA^11^. The dichotomizing process combined multiple levels into one group, which could lose useful phenotypic information and statistical power. On the other hand, treating the categorical phenotypes as a quantitative trait violates the nature of the ordinal categorical phenotype, which could result in inflated type I error rates and power loss. Through simulation studies and real data analysis, unless the phenotypic distribution is extremely unbalanced, the linear mixed model approaches are still reliable when testing common variants, which suggests that fastGWA analyses that limited to SNPs with MAF > 0.01 (http://fastgwa.info/ukbimp/phenotypes) should still be valid for many of the phenotypes. However, considering the diversity of the phenotypic distribution, it is difficult to select a MAF cutoff to remain the association testing accurate in practical. In addition, we identified many phenotypes associated variants with MAF < 0.01 in the UK-Biobank data analysis, that were missed in the fastGWA analyses.

We applied the FastPOLMM to analyze 258 ordinal categorical phenotypes on UK Biobank, of which 150 phenotypes are food and other preferences (UK Biobank Category 1039). The preference data (version 1.1) was released in January 2020. To the best of our knowledge, this is the first time that GWAS were applied to analyze the preference data. All analyses results have been made publicly available through a visual server (http://polmm.leelabsg.org/). The web interface provides intuitive visualizations and is useful resource for post-GWAS analyses.

There are several limitations in POLMM, most of which are similar as SAIGE and other mixed model approaches. First, DensePOLMM is still computationally expensive when fitting a null mixed model with greater than 500,000 samples. Second, POLMM methods estimate odds ratios for genetic markers using the parameter estimates from the null model and might not be accurate. Third, POLMM assumes an infinitesimal architecture, that is, the effect sizes of genetic markers are normally distributed. If the genetic architecture is non-infinitesimal, POLMM methods may sacrifice power. Finally, the variance component estimate 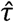 is biased and should not be used to estimate heritability. Interestingly, we observe a more accurate estimate 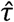 as the number of categories increases. Although based on the proportional odds assumption, POLMM approaches can still control type I error rates at a stringent significance level of 5 × 10^−8^ when categorical phenotypes follow other models including adjacent category logistic model and stereotype model (Figure S16).

In the future, we plan to extend the current single-variant test to gene-or region-based multiple variants tests to better identify the rare variants. Recently, a novel machine learning method called REGENIE was proposed for quantitative and binary traits analysis. Instead of using mixed effect model, REGENIE^26^ uses ridge regression model to account for polygenic effects. We plan to evaluate the strategies in REGENIE in ordinal categorical data analysis to extend POLMM. POLMM approaches are motivated to analyze large-scale biobank data collected following a cohort study design. Suppose that data is collected from a matched case-control study design, the stratified sampling for different levels could inflate the parameter estimation and genetic association testing.^27^ We plan to extend the POLMM approaches to deal with the effect of the sampling.

Ordinal categorical phenotypes are widely observed in survey, questionnaires, and testing to measure human behaviors, satisfaction, and preferences. However, due to the lack of analysis tools, methods designed for binary and quantitative traits have been used to analyze the categorical data, which is inappropriate and can result in suspicious results. Our method POLMM provides an accurate and scalable solution with the following features: can accurately model the ordinal categorical data using a proportional odds logistic model which can; can adjust for sample relatedness by incorporating random effects; can be scalable to analyze a large-scale dataset with hundreds of thousands of subjects; can test low-frequency variants under unbalanced phenotypic distribution by using SPA to approximate the null distribution of the test statistics. Due to all these features, POLMM is a unified and the only available approach for ordinal categorical data analysis in biobanks and large cohort studies.

### URLs

POLMM (version 0.2.2), https://github.com/WenjianBI/POLMM.

BOLT-LMM (version 2.3.4), https://alkesgroup.broadinstitute.org/BOLT-LMM

SAIGE (version 0.36.3), https://github.com/weizhouUMICH/SAIGE.

fastGWA (GCTA, version 1.93.1beta), https://cnsgenomics.Com/software/gcta/#fastGWA.

UK Biobank PheWeb and analysis results, http://polmm.leelabsg.org/.

ANNOVAR (16 Apr 2018), https://doc-openbio.readthedocs.io/projects/annovar/en/latest/

## Acknowledgements

This research was supported by NIH grants R01-HG008773 (W.B. and S.L.), and Brain Pool Plus (BP+, Brain Pool+) Program through the National Research Foundation of Korea (NRF) funded by the Ministry of Science and ICT (2020H1D3A2A03100666, S.L). UK Biobank data was accessed under the accession number: 45227.

## Author Contributions

W.B., J.N.S, and S.L. designed the experiments. W.B. and S.L. developed the software tools, performed the simulations, and analyzed the UK Biobank data under the assistance and guidance from W.Z., R.D., B.M, and J.N.S. W.B. and S.L. wrote the manuscript with the participation of all authors. All authors reviewed and approved the final manuscript.

## Competing Interests statement

The authors declare no competing interests.

## Methods

### Data Simulation

In simulation studies, genotypes were simulated based on the pedigree shown in Figure S1, in which each family includes 10 subjects. To estimate GRM for mixed models fitting, we simulated 100,000 independent SNPs with MAFs ranging from 0.05 to 0.5. For subject *i*, two covariates *X*_*i*1_ and *X*_*i*2_ were simulated following the standard normal distribution and a Bernoulli (0.5) distribution, respectively. Given the variance component *τ*, random effects *b = (b*_1_, *b*_2_, …, *b*_*n*_) were simulated following a multivariate normal distribution *N*(0, *τV*) where *V* is the GRM from the family structure. We followed model (1) to simulate ordinal categorical phenotypes using linear predicator *η*_*i*_ *=* **0**.**5** · *X*_*i*l_ **+ 0**.**5** · *X*_*i*2_ *+ γ*· *G*_*i*_ *+ b*_*i*_, *i* ≤ *n*, in which *G*_*i*_ is the genotype value of one SNP. We considered two common types of phenotypic distribution: bell-shaped distribution with three categories and L-shaped distribution with four categories (Figure S3), and selected cutpoints *ϵ* to correspond to the given phenotypic distribution.

Under the null model *γ =* **0**, we considered three variance components *τ* =0.5, 1, and 2 to evaluate type I error rates at a significance level *α =* 5 × 10^−8^. For each phenotypic distribution, we simulated 100 datasets of phenotypes. We considered common, low-frequency, and rare SNPs with MAFs of 0.3, 0.01, and 0.005, respectively. For each MAF, we simulated 10^7^ SNPs. Thus, for each pair of phenotypic distribution and MAF, totally 10^9^ tests were performed. Under the alternative model *γ* ≠ 0, we considered variance component *τ* = 1 and increased genetic effect size *γ* to evaluate empirical powers at a significance level *α =* 5 × 10^−8^. For each *γ*, we simulated 200 datasets including ordinal categorical phenotypes and genotypes of one causal SNP. Since BOLT-LMM methods cannot control type I error rates in some scenarios, we used empirical significance levels to calculate the empirical powers.

To compare DensePOLMM and FastPOLMM, we added a scenario to simulate random effect vector *b*. First, we randomly selected 50,000 SNPs (i.e. 50%) from the 100,000 SNPs that were used to estimate GRM. Then, for subject *i*, random effect 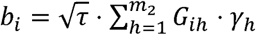 where *m*_2_ *=* **50**,**000**, *G*_*ih*_ was the genotype of the *h*-th selected SNP, and *γ*_*h*_ was simulated following a normal distribution *N*(0,0.085) so that the empirical variance of the random effects is close to *τ*. In this scenario, the random effects were strongly related to the estimated GRM used in the null mixed models fitting. We set variance components *τ* = 1 and 10 to simulate moderate and high heritability, respectively. We simulated phenotypes with 5 and 10 evenly distributed categories.

### Details for Runtime Evaluation

All analyses were conducted on CPU cores of Intel(R) Xeon(R) Gold 6138 @ 2.00GHz. In step 1, we used 8 CPU cores and recorded the computation time. For SAIGE, fastGWA, and POLMM methods, the null mixed model fitting result can be saved and used for association testing. Hence, the genotype data to test can be divided into multiple chunks for parallel computation. In step 2, we used 1 CPU core and recorded the computation time. For BOLT-LMM, the model fitting and association testing cannot be separately implemented. We extracted “the time for streaming genotypes and writing output” from log files to record the computation time in step 2. Since FastPOLMM and DensePOLMM are the same when testing genetic association effect, we only recorded the computation time of DensePOLMM in step 2.

### Liability Threshold Model and Liability Heritability

Model (1) is equivalent to the following liability threshold model

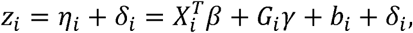

where *z*_*i*_ is a latent variable and error term *δ*_*i*_ follows a logistic distribution with a location parameter of 0 and a scale parameter of 1. The ordinal categorical phenotype *y*_*i*_ *= j* if the latent variable *z*_*i*_ is between cutpoints *ϵ*_*j −* 1_ and *ϵ*_*j*_. The variances of *b*_*i*_ and *δ*_*i*_ are *τ* and π^2^/3, respectively. Hence, similar to SAIGE^8^, we define a liability heritability 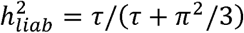. Variance components *τ* = 1 and 10 correspond to liability heritability 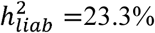 and 75.2%, respectively.

### Maximum likelihood estimation and score test

For mathematical convenience, we define a *J* × 1 vector 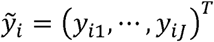 as an equivalent representation of the ordinal categorical phenotype *y*_*i*_ : if *y*_*i*_ *= j*, then *y*_*ij*_ *=* 1 and the other elements in 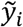 are 0. For subject *i*, the log-likelihood function given random effects *b* is

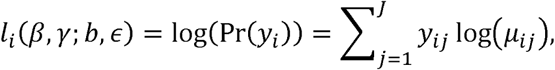

where *μ*_*ij*_ is the mean of *y*_*ij*_, that is,

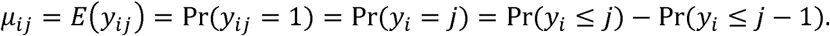

Similar to SAIGE^8^, we propose an efficient algorithm (Supplementary Note) to iteratively estimate parameters using PQL and AI-REML algorithm to maximize log-likelihood function

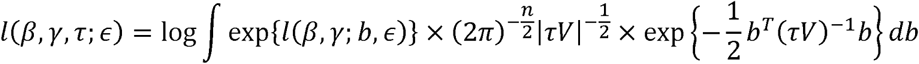

where *l*(*β, γ*; *b, ϵ*) = Σ_*i*≤*n*_ *l*_*i*_ (*β, γb, ϵ*), and then use a hybrid strategy of normal approximation and SPA to calculate p values.

### Conditional analysis

We let *G* denote the n-dimensional genotype vector of the marker to test and let *G*_*c*_ denote the n-dimensional genotype vector of the conditioning marker. The covariate-adjusted genotype vectors

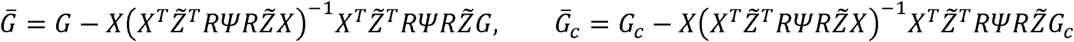

correspond to the two genotype vectors. The definitions of matrix *X*, 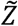, *R, W*, and Pcan be seen in Supplementary Note. Under the null hypothesis, the conditional score statistic is

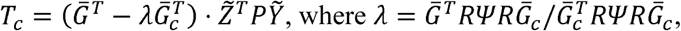

and its estimated variance 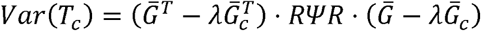. Then, we can use similar hybrid strategies to test the conditional score statistic using normal distribution approximation and SPA.

### Parameter estimation

Fitting an alternative mixed model is required to accurately estimate the parameter 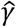 and the corresponding odds ratio 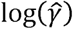. However, it takes much time and is not scalable for a genome-wide association study. We used similar strategy as in SAIGE to use the information from the null model fitting to estimate the parameter

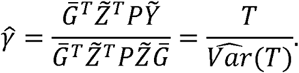

Since both *T* and 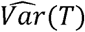 have been calculated for association p-value estimation, it does not require additional computations. We use p values to estimate the standard error of 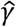 as 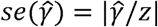, where z-score corresponds to the association p value / 2.

### Leave-one-chromosome-out scheme

To avoid contamination for correlated markers, we implemented an option to apply the leave-one-chromosome-out (LOCO) for DensePOLMM and FastPOLMM methods. If LOCO scheme is used, we first use all SNPs to estimate the variance component 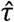, and then for each chromosome, we updated the estimation of 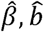, and 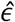 after excluding all SNPs in the same chromosome. This strategy is the same as SAIGE and BOLT-LMM. For FastPOLMM, we first used tool GCTA to calculate GRM for each chromosome and then combined them to calculate GRMs.

### Approaches to Reducing Computation Time and Memory Cost

To make DensePOLMM method computationally practical for studies with large sample size *n*, we use strategies as in BOLT-LMM^10^ and SAIGE^8^ to reduce computational and memory cost. Instead of storing an *n*×*n* dimensional dense GRM, we compactly store raw genotypes of the genetic variants into a binary vector and use them when dense GRM is needed. When fitting the null mixed model and estimating variance 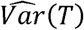, we need solve linear system Σ · *x = u*, which is challenging since Cholesky decomposition takes *O*(*n*^3^) computation and very large memory space to invert matrix Σ. For a given vector *u*, we use PCG approach^8^ to directly calculate Σ^−1^*u*. To make the convergence faster, we use a block diagonal matrix *Q =* diag(*Q*_1_, …, *Q*_*n*_) as the preconditioner matrix, where (*J* − 1) × (*J* − 1) matrix 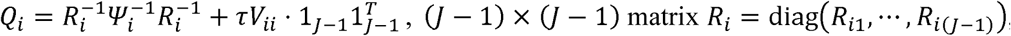, (*J* − 1) × (*J* −1) matrix *R*_*i*_ *=* diag(R_*i*1_,…,*R*_*i*(*J*−1)_), and (*J* −1) dimensional vector of ones l_*j*−1_ = (1,1, …,1)^*T*^. Given the same tolerance criterion as in SAIGE, PCG in POLMM usually takes 6-8 iterations to converge, which is ∼ 1.5 times more than that in SAIGE. This might be because that we use a block diagonal matrix as the preconditioner matrix, in which each block corresponds to one subject. When updating variance component 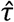, we estimate 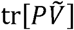 by using Hutchinson’s randomized trace estimator, 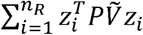 where 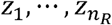 are *n*_*R*_ independent random vectors whose elements are i.i.d Rademacher random variables.^28^ In addition, we use Intel Threading Building Block (TBB) implemented in RcppParallel package^29^ for the multi-threading computation.

### Genome build

All genomic coordinates are given according to NCBI Build 37/UCSC hg19.

---

**Figure S1.**
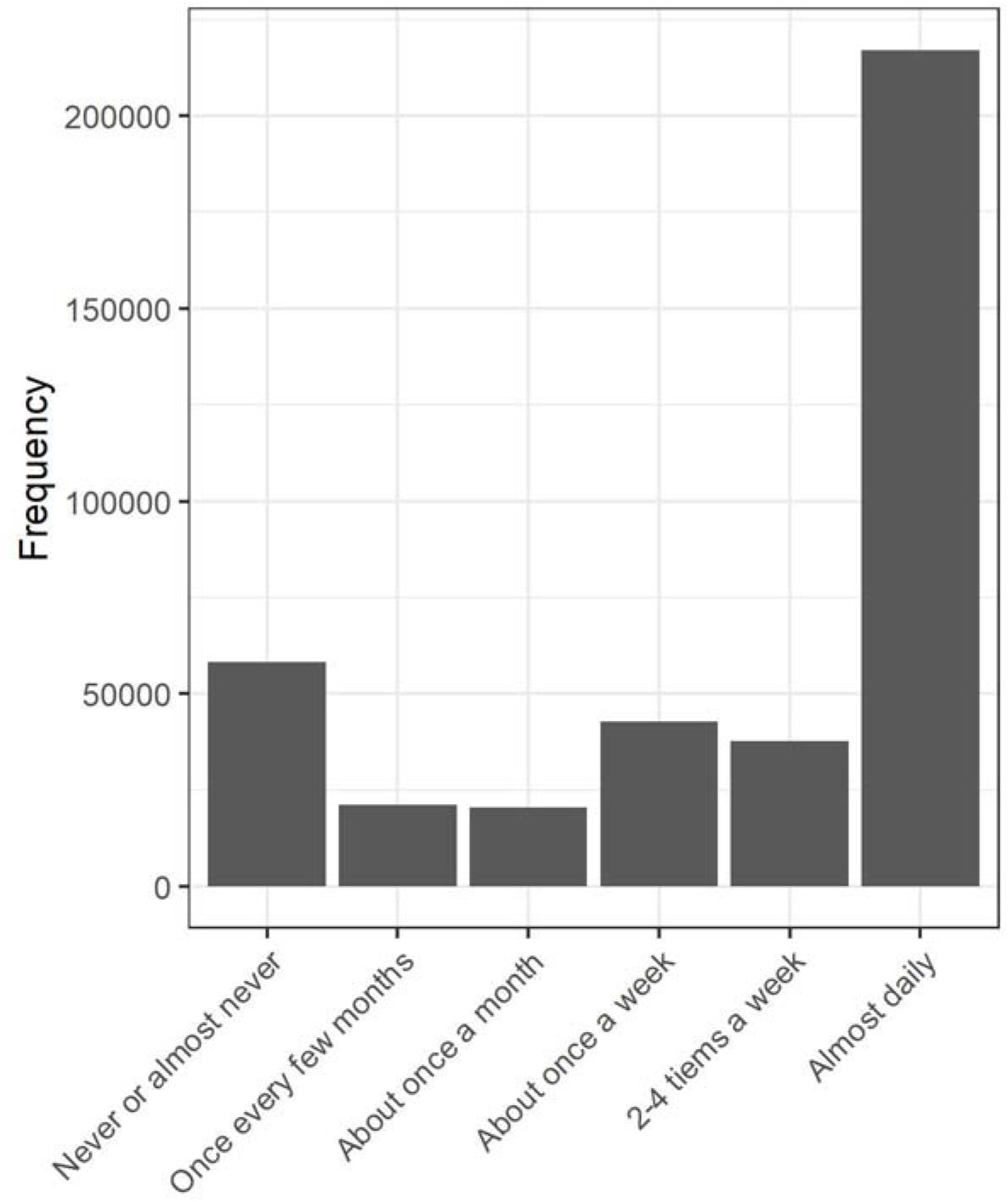
Sample size distribution of categorical phenotype, able to confide, in UK Biobank. For SAIGE, we defined a binary phenotype as 1 or 0 depending on whether the categorical phenotype is “almost daily” or not. For POLMM methods, we combined subjects that are neither “never or almost never” nor “almost daily” to make a categorical phenotype with 3 levels.

**Figure S2.**
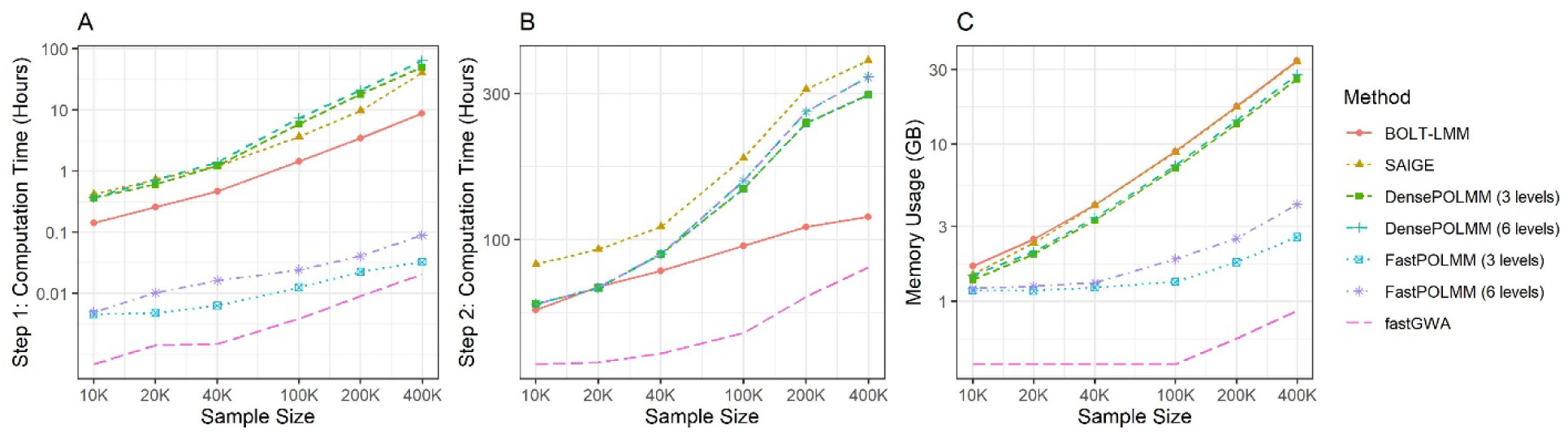
Computation time and memory usage of BOLT-LMM, SAIGE, DensePOLMM, FastPOLMM, and fastGWA. CPU core is Intel(R) Xeon(R) Gold 6138 @ 2.00GHz. (A). Computation time in step 1 to fit a null mixed model; (B) Computation in step 2 to test 30 million variants; (C) Memory usage in step 1, fastGWA requires less than 0.4 GB memory when sample size is less than 100,000.

**Figure S3.**
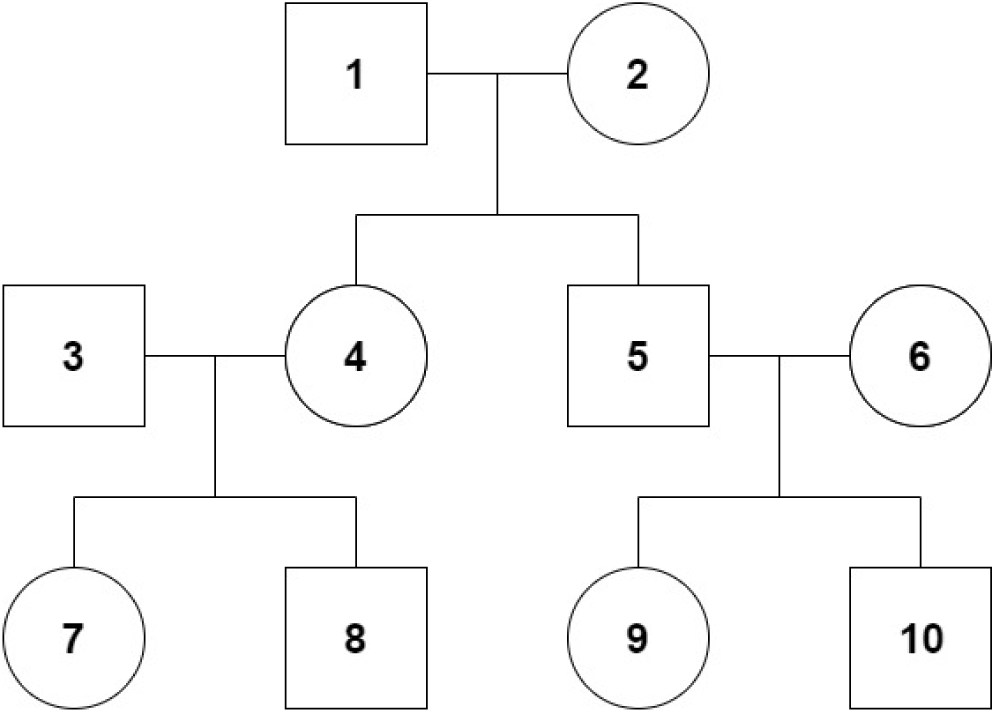
Pedigree of families in simulation studies

**Figure S4.**
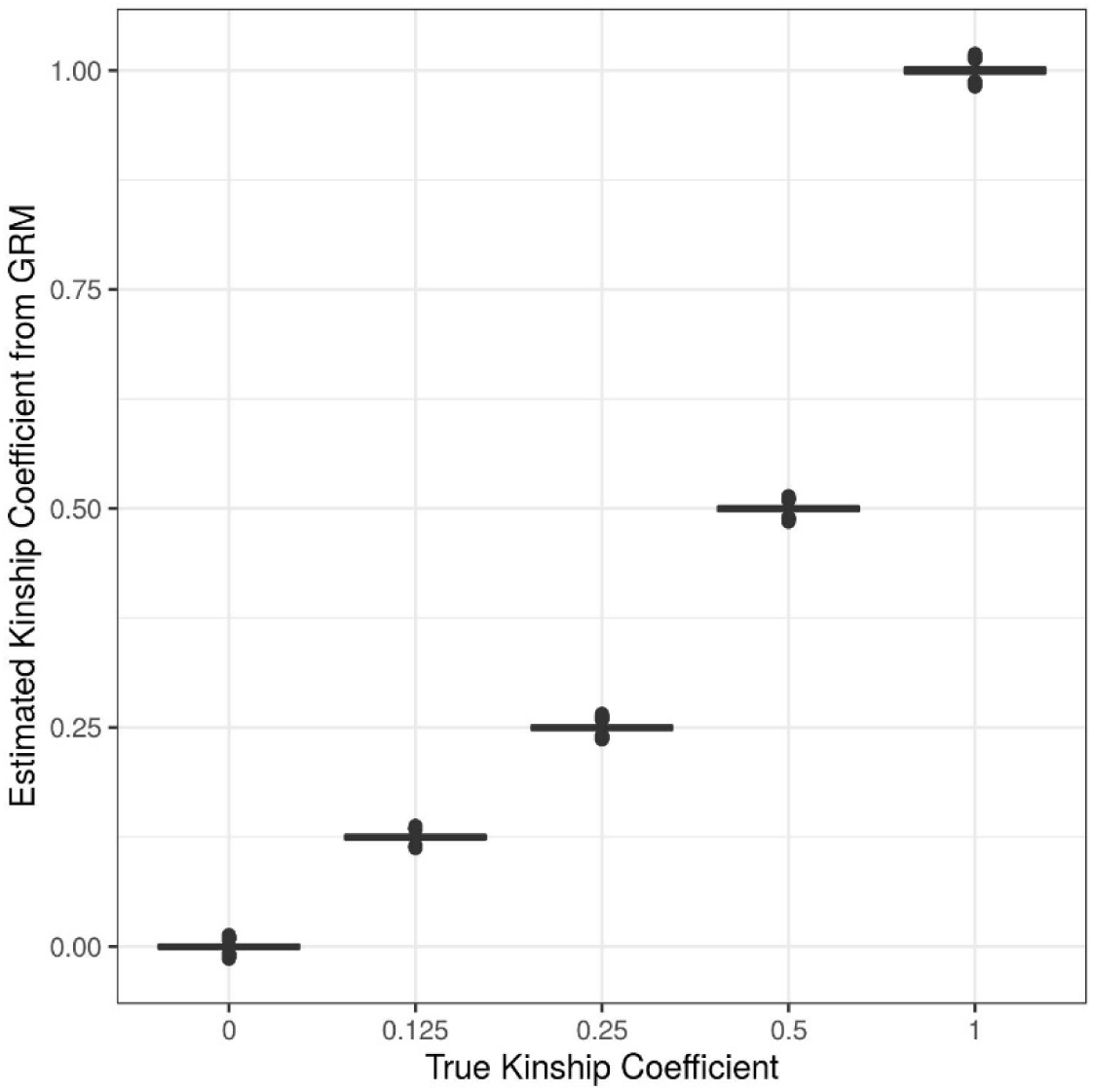
Kinship coefficients estimated from the empirical GRM in simulation studies

**Figure S5.**
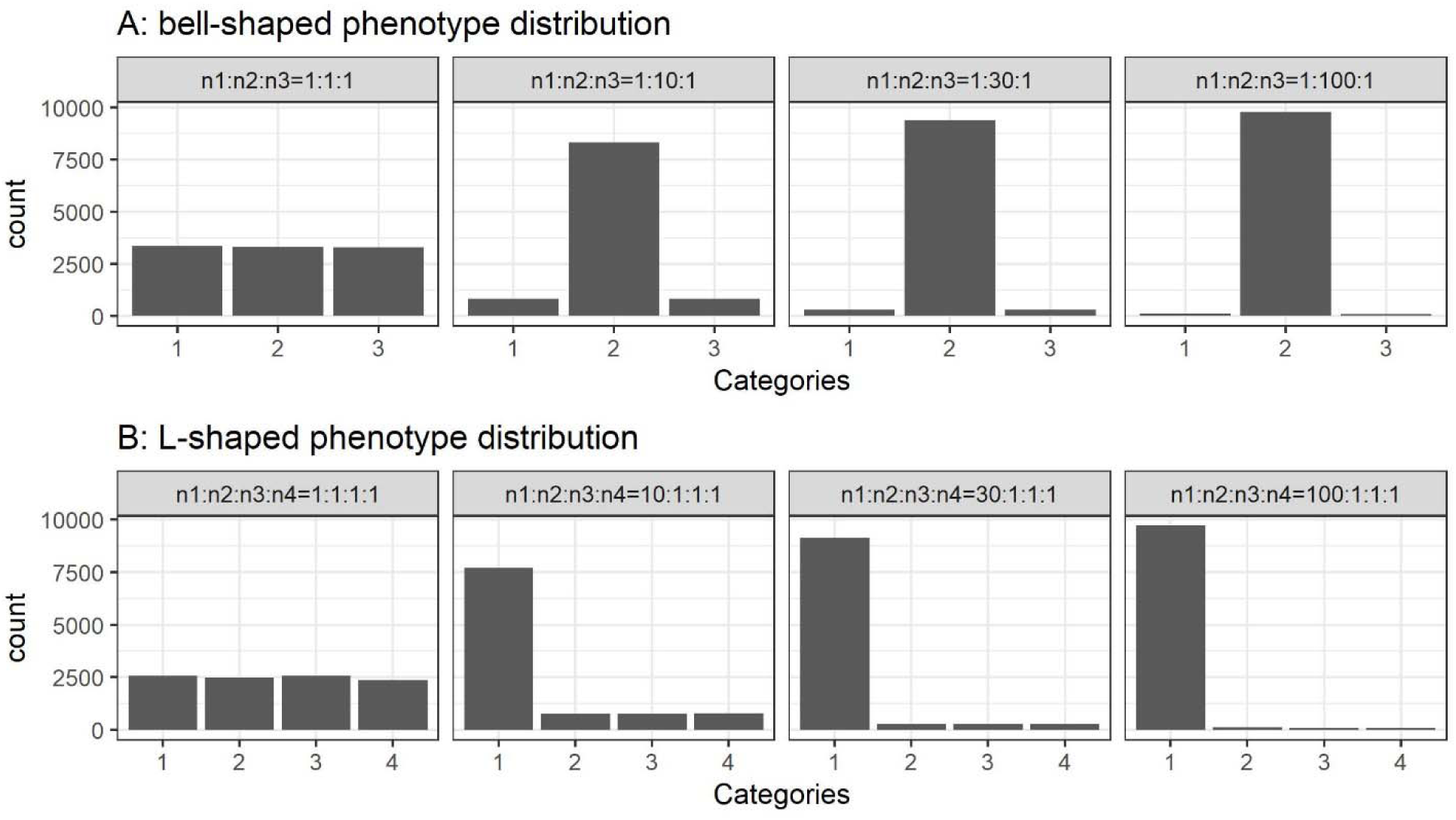
Sample size distribution of the ordinal categorical phenotypes in simulation studies. We simulated 1,000 families with a total sample size *n =* 10,000. (A) Bell-shaped phenotype distribution with three levels. The sample sizes in three levels are *n*_1;_ *n*_2_, and *n*_3_, respectively. From left to right, we simulated four scenarios: balanced (*n*_1_: *n*_2_:*n*_3_ *=* 1:1:1), moderately unbalanced (*n*_1_: *n*_2_:*n*_3_ *=* 1:10:1), unbalanced (*n*_1_: *n*_2_:*n*_3_, = 1:30:1), and extremely unbalanced (*n*_1_: *n*_2_:*n*_3_, = 1:100:1). (B) L-shaped phenotype distribution with four levels. The sample sizes in four levels are *n*_1_, *n*_2_, *n*_3_, and *n*_4_, respectively. From left to right, we simulate four scenarios: balanced (*n*_1_: *n*_2_:*n*_3_:*n*_4_ *=* 1:1:1:1), moderately unbalanced (*n*_1_ *n*_2_: *n*_3_: *n*_4_ = 10:1:1:1), unbalanced (*n*_1_. *n*_*2*_: *n*_*3*_: *n*_*4*_ *=* 30:1:1:1), and extremely unbalanced (*n*_1_. *n*_*2*_: *n*_*3*_:*n*_*4*_= 100:1:1:1). For bell-shaped phenotype distribution, we use two methods to dichotomize the categorical phenotype to evaluate SAIGE. SAIGE-1: level 1 versus levels 2 and 3; SAIGE-2: levels 1 and 2 versus level 3. For L-shaped phenotype distribution, we dichotomize the phenotype: level 1 versus levels 2-4.

**Figure S6.**
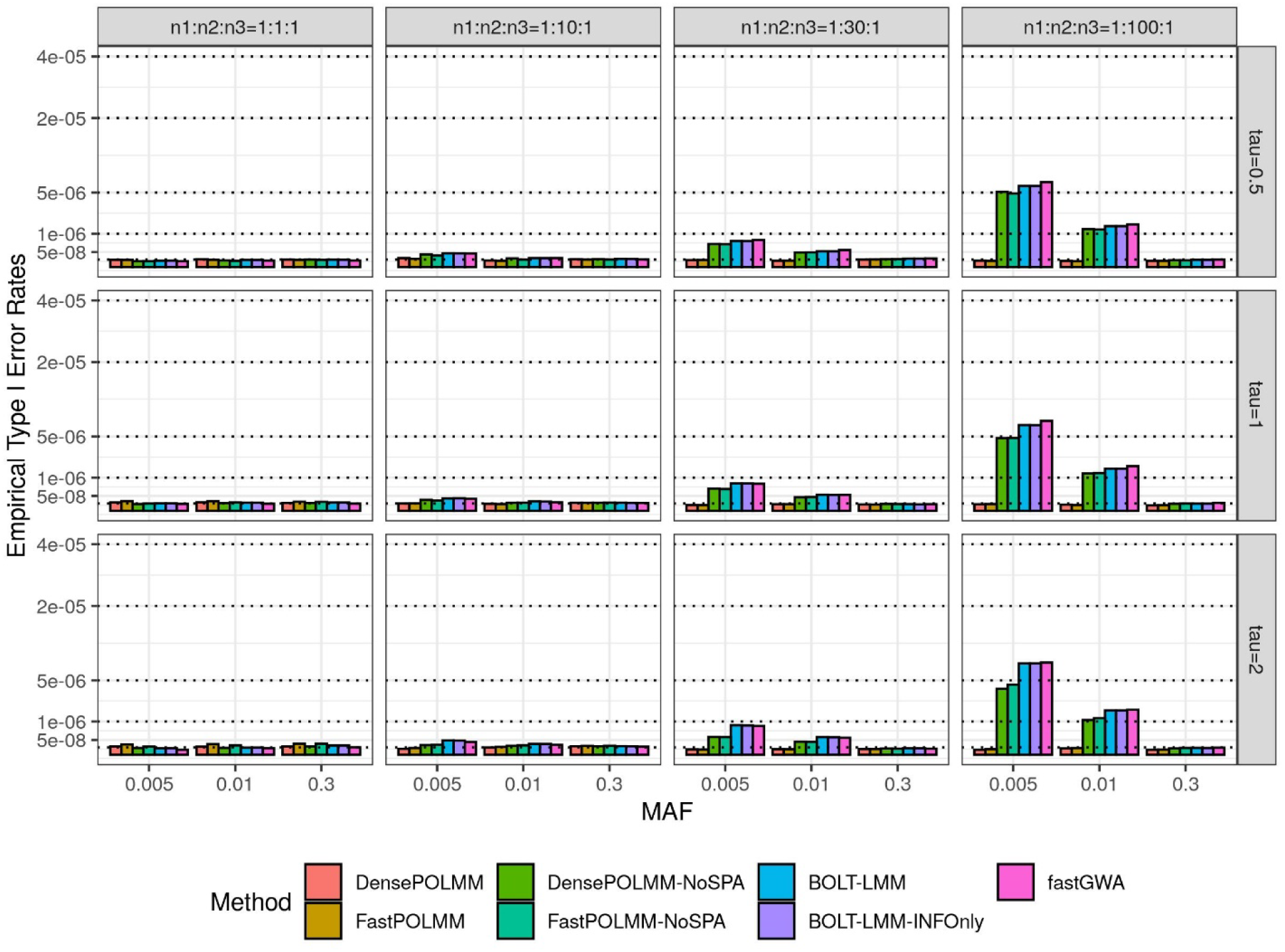
Empirical type I error rates of POLMM, BOLT-LMM, and fastGWA methods at a significance level 5e-8. We simulated 1,000 families with a total sample size *n =* 10,000 and an ordinal categorical phenotype including three levels with sample sizes *n*_1;_ *n*_2_, and *n*_3_. From left to right, the plots consider four scenarios: balanced (*n*_1_: *n*_2_:*n*_3_ = 1:1:1), moderately unbalanced (*n*_1_: *n*_2_:*n*_3_ = 1:10:1), unbalanced (*n*_1_: *n*_2_:*n*_3_ *=* 1: 30:1), and extremely unbalanced (*n*_1_: *n*_2_:*n*_3_ = 1:100:1). From top to bottom, the plots consider three variance components *τ*= 0.5, 1, and 2. We simulated common, low-frequency, and rare variants with MAFs of 0.3, 0.01 and 0.005, respectively.

**Figure S7.**
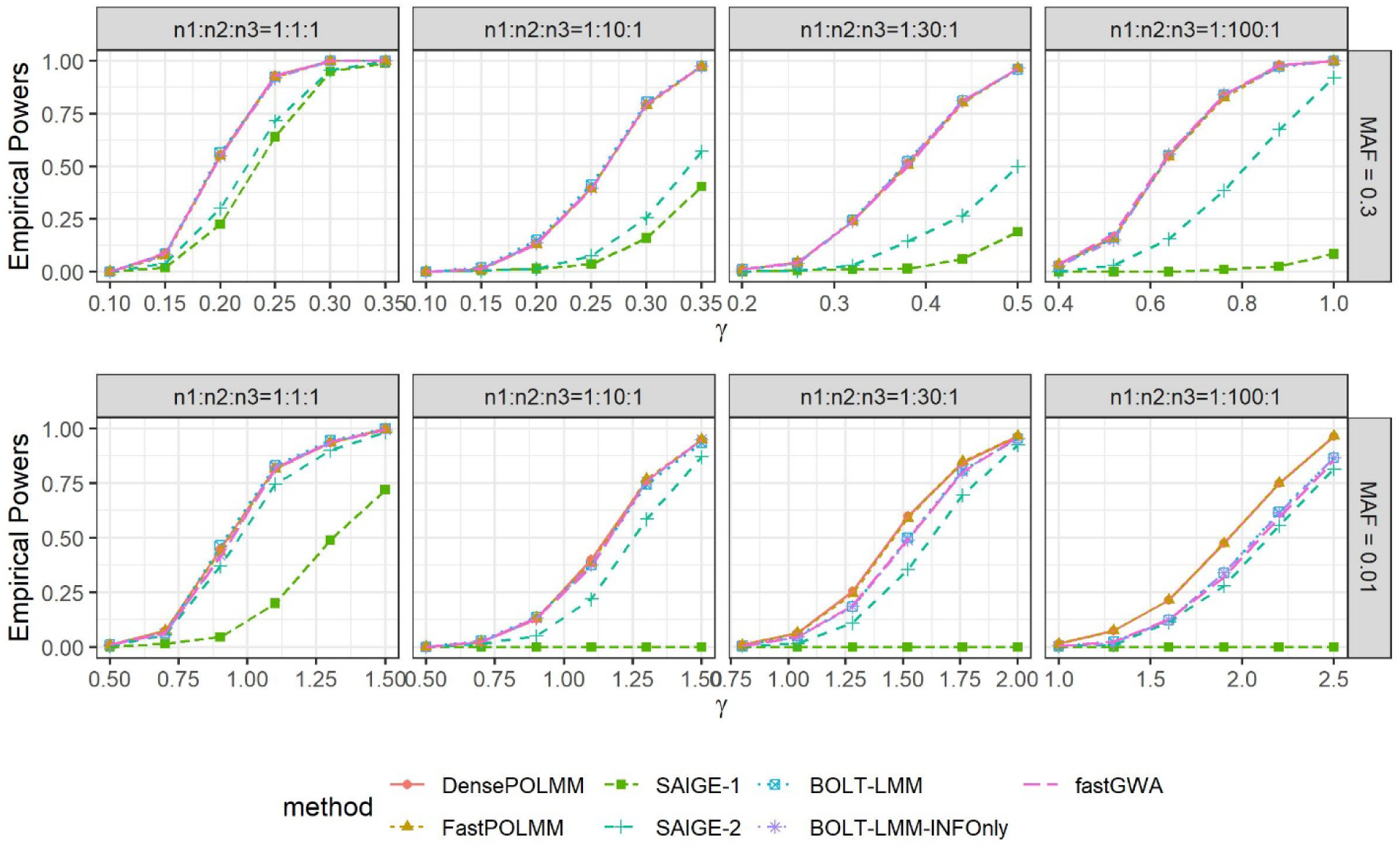
Empirical powers of DensePOLMM, FastPOLMM, SAIGE, BOLT-LMM, and fastGWA methods at a significance level 5e-8. We simulated 1,000 families with a total sample size *n =* 10,000 and an ordinal categorical phenotype including three levels with sample sizes *n*_1;_ *n*_2_, and *n*_3_. From left to right, the plots consider four scenarios: balanced (*n*_1_: *n*_2_:*n*_3_ = 1:1:1), moderately unbalanced (*n*_1_: *n*_2_:*n*_3_ = 1:10:1), unbalanced (*n*_1_: *n*_2_:*n*_3_ = 1: 30:1), and extremely unbalanced (*n*_1_: *n*_2_:*n*_3_ = 1:100:1). From top to bottom, the plots consider two MAFs of 0.3 and 0.01 to simulate common and low-frequency variants. We let variance component *τ* = 1. For SAIGE, we use different cutoffs to dichotomize phenotypes (SAIGE-1: level 1 as controls and levels 2,3 as cases; SAIGE-2: levels 1,2 as controls and level 3 as cases). For BOLT-LMM, the empirical powers were calculated based on the empirical significance levels since it cannot control type I error rates for low-frequency variants.

**Figure S8.**
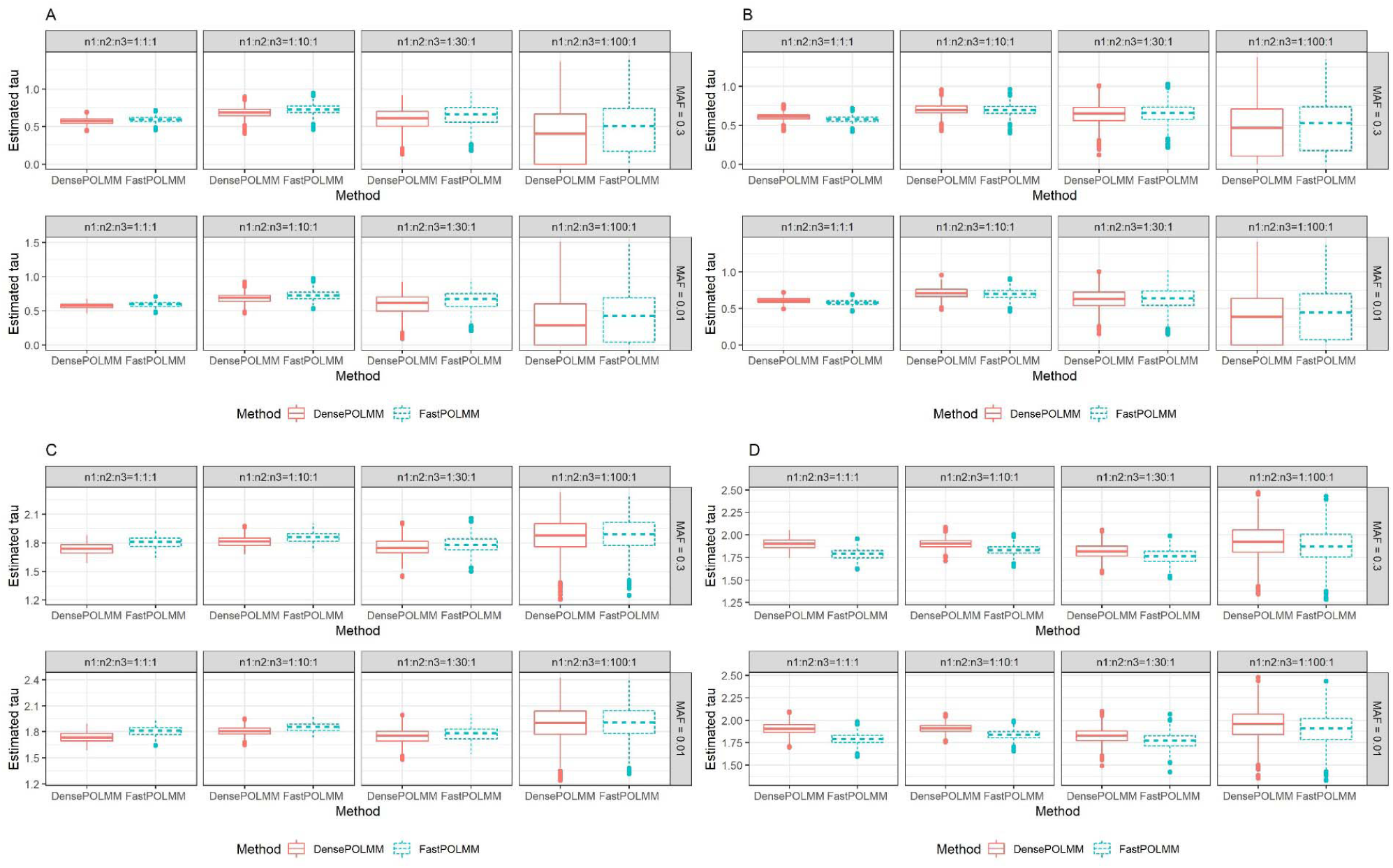
Estimated variance component of DensePOLMM and FastPOLMM when number of levels is 3. (A) vector *b* is simulated following *N*(0, *τV)* and *τ* = 1; (B) vector *b* is simulated based on the SNPs used in GRM and *τ* = 1; (C) vector *b* is simulated following *N(0, τV)* and *τ*= 10; (D) vector *b* is simulated based on the SNPs used in GRM and *τ*= 10.

**Figure S9.**
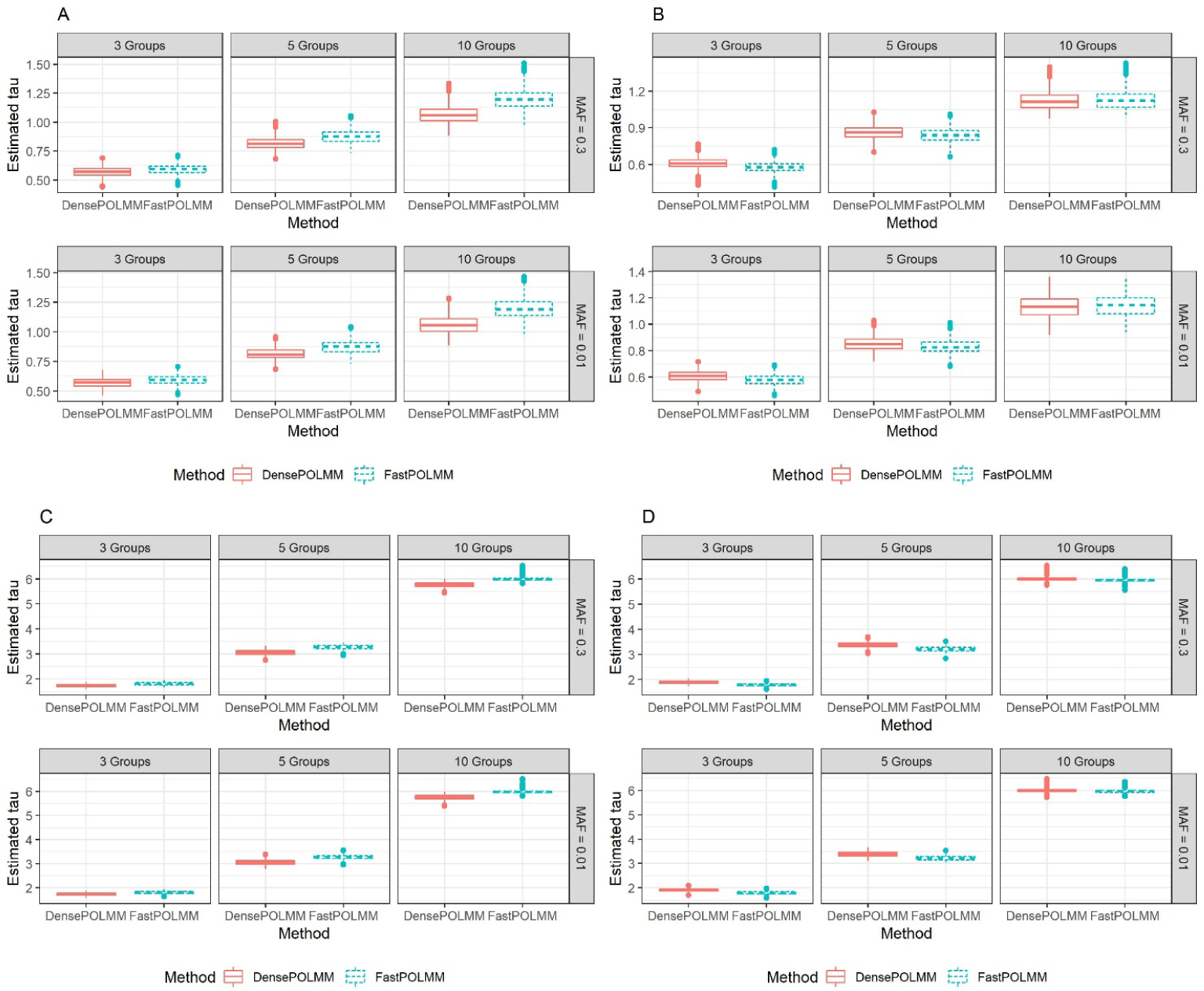
Estimated variance component of DensePOLMM and FastPOLMM when the number of evenly distributed levels is 3, 5, and 10. (A) vector *b* is simulated following *N*(0, *τV)* and *τ* = 1; (B) vector *b* is simulated based on the SNPs used in GRM and *τ* = 1; (C) vector *b* is simulated following *N(0, τV)* and *τ*= 10; (D) vector *b* is simulated based on the SNPs used in GRM and *τ*= 10.

**Figure S10.**
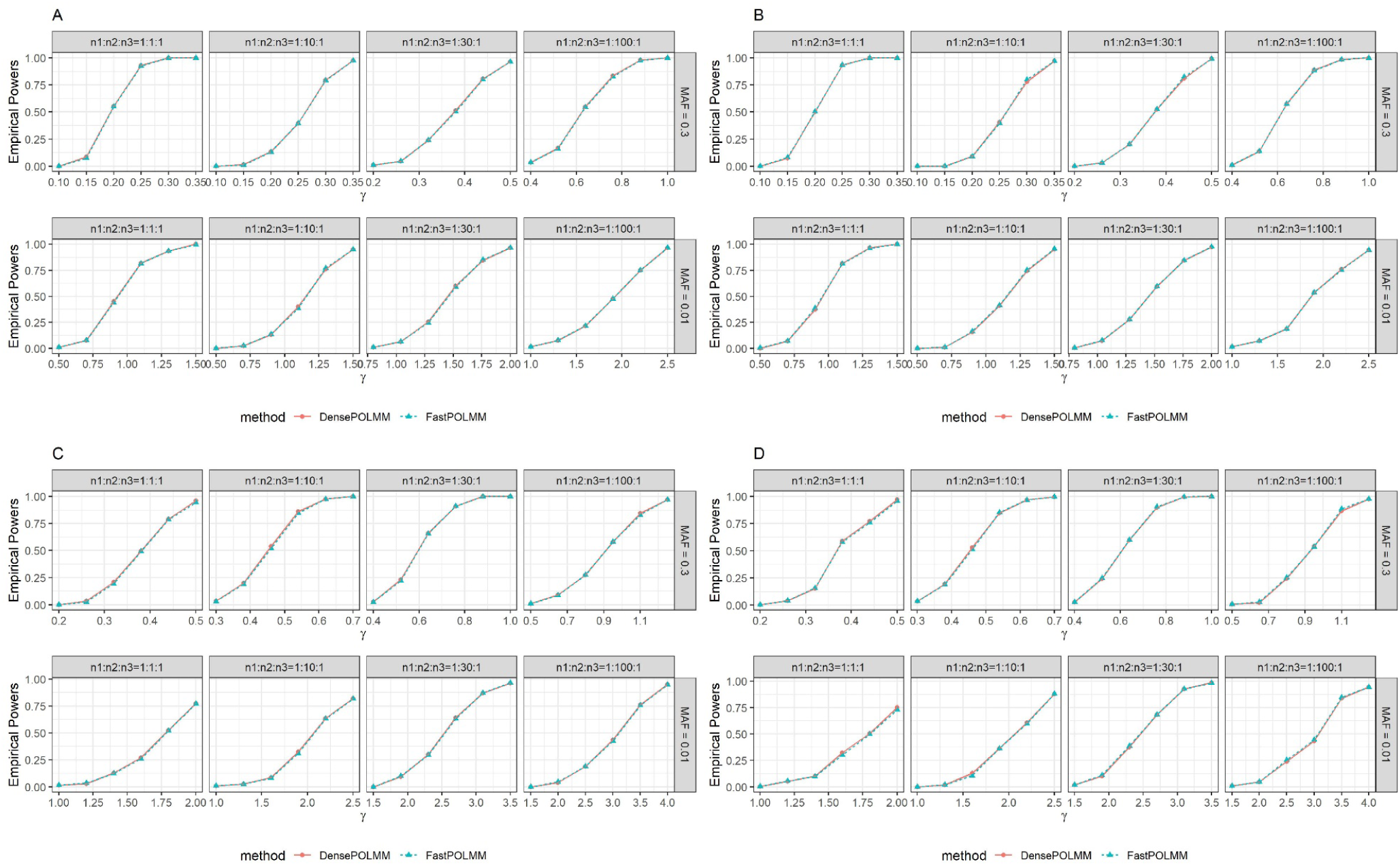
Empirical powers of DensePOLMM and FastPOLMM when the number of levels is 3. (A) vector *b* is simulated following *N(Q, τV)* and *τ = l;* (B) vector *b* is simulated based on the SNPs used in GRM and *τ*= 1; (C) vector *b* is simulated following *N(0, τV)* and *τ* = 10; (D) vector *b* is simulated based on the SNPs used in GRM and *τ* = 10.

**Figure S11.**
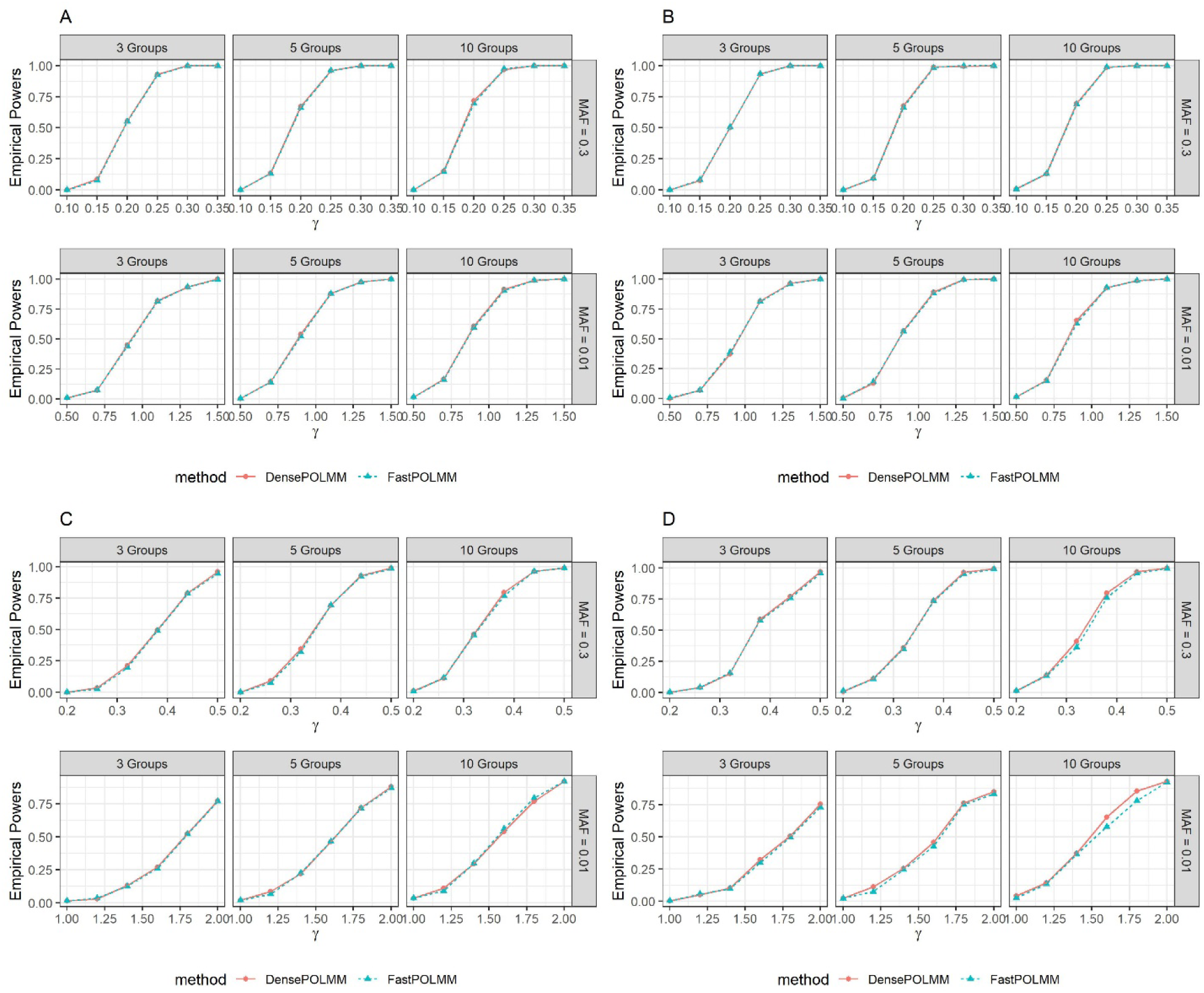
Empirical powers of DensePOLMM and FastPOLMM when the number of evenly distributed levels is 3, 5, and 10. (A) vector *b* is simulated following *N*(0, *τV*) and *τ* = 1; (B) vector *b* is simulated based on the SNPs used in GRM and *τ*= 1; (C) vector *b* is simulated following *N*(0, *τV*) and *τ*= 10; (D) vector *b* is simulated based on the SNPs used in GRM and *τ* = 10.

**Figure S12.**
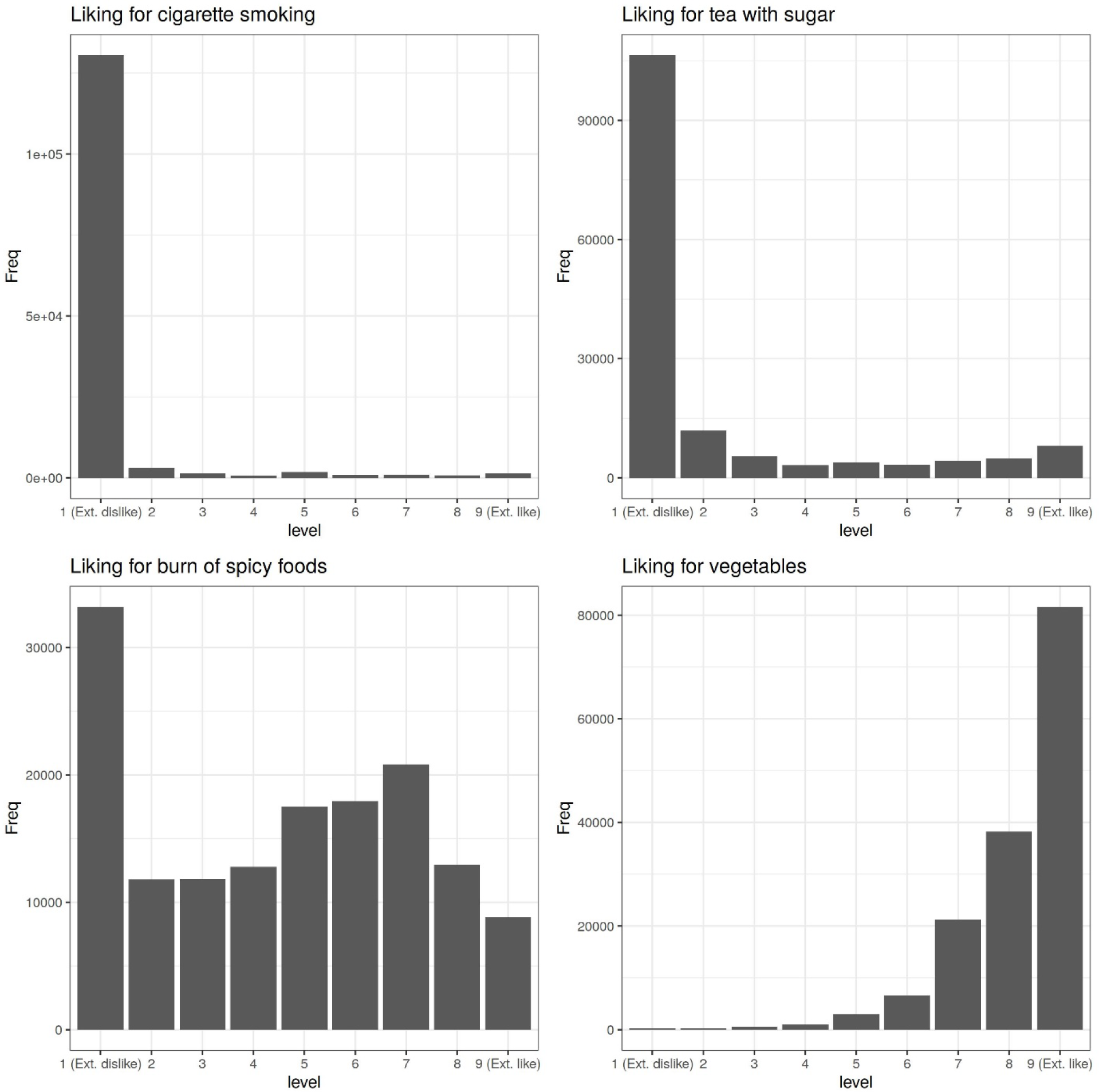
Sample size distribution of the four ordinal categorical phenotypes selected to compare POLMM and BOLT-LMM methods in UK Biobank data analysis

**Figure S13.**
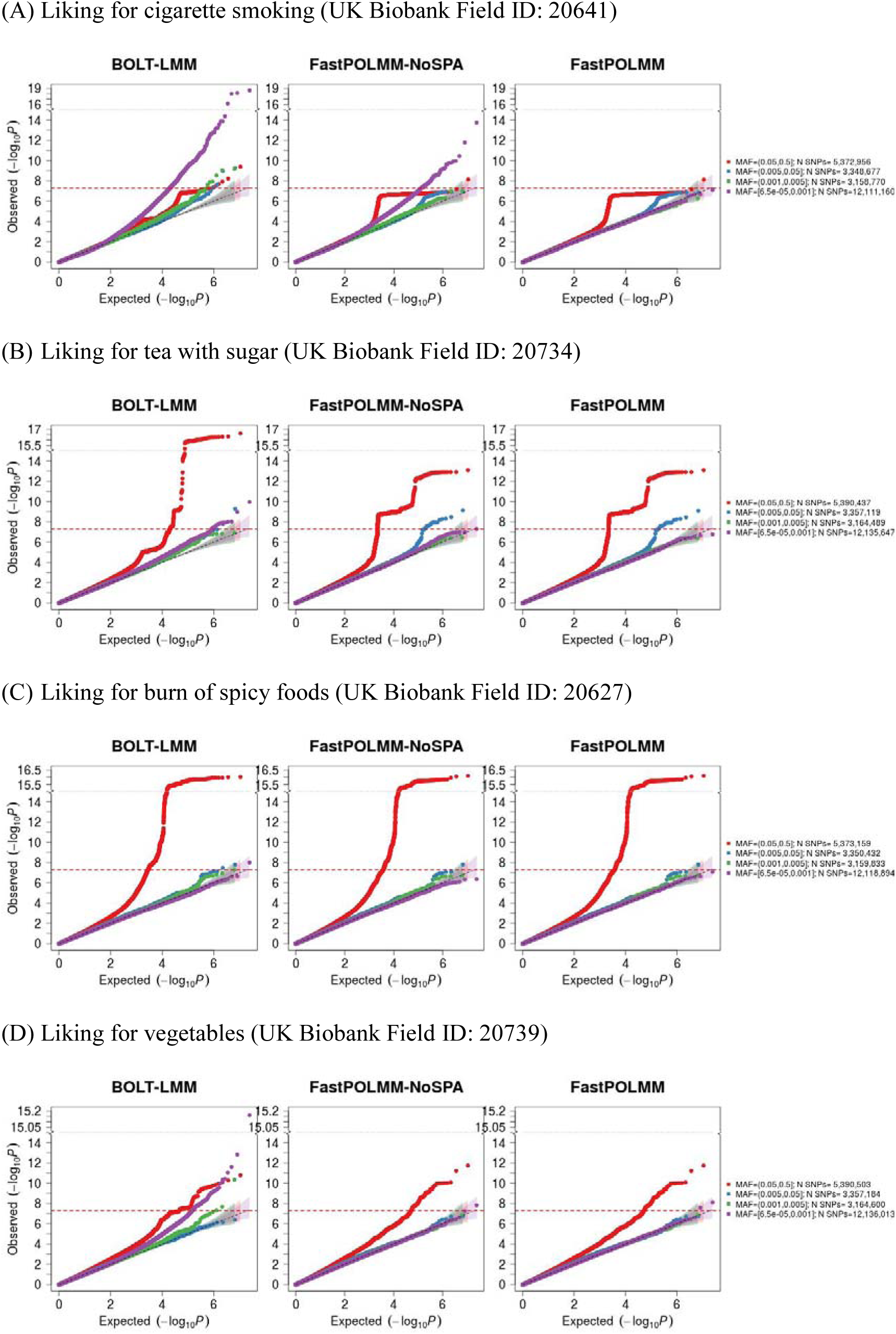
QQ plots for UK Biobank analysis

**Figure S14.**
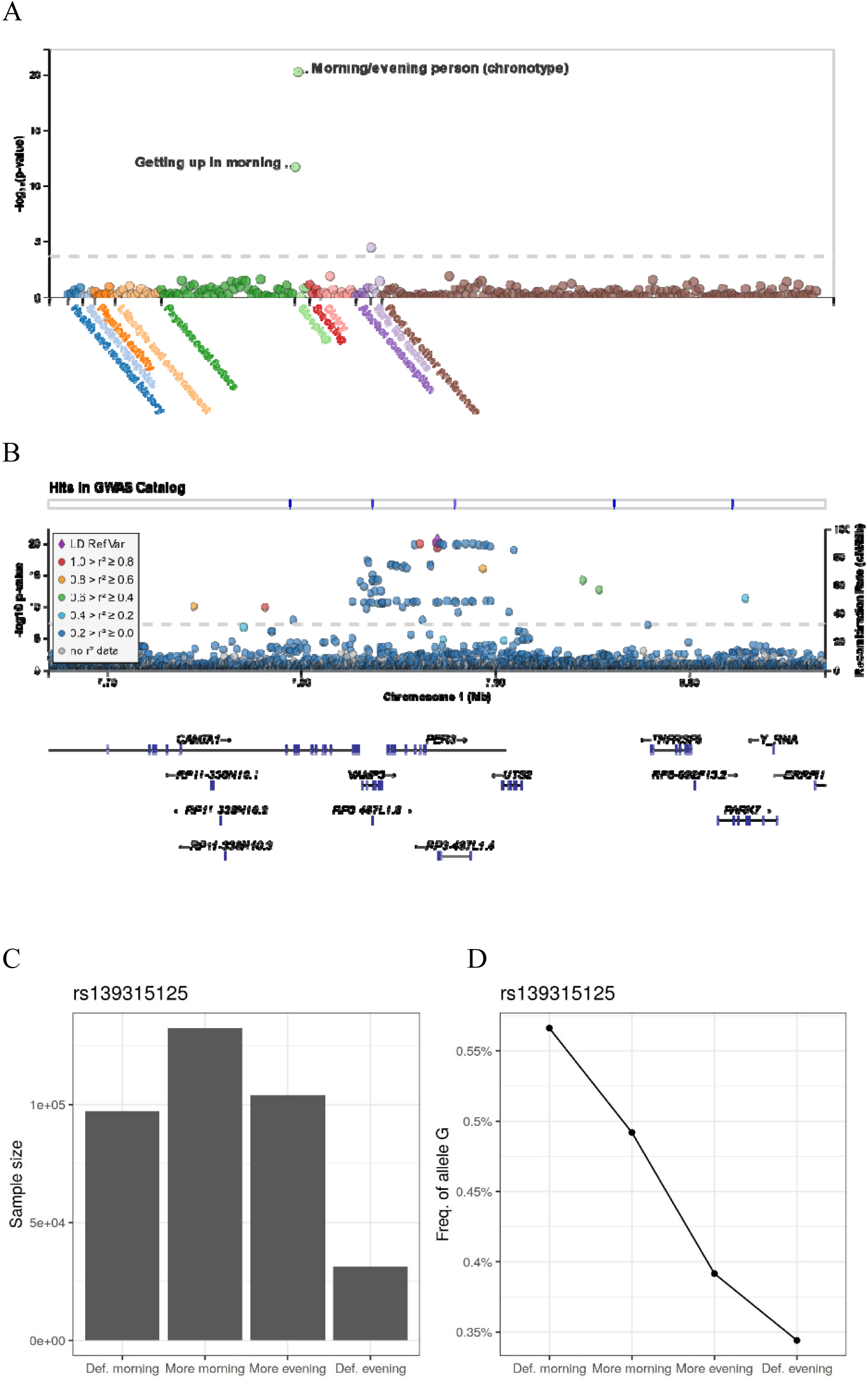
Morning/evening person versus rs139315125, a nonsynonymous single nucleotide variant in PER3 Gene. The phenotypes (Field ID: 1180) include 4 ordinal categories: definitely an “evening” person, more an “evening” than a “morning” person, more a “morning” than “evening” person, and definitely a “morning” person. (A) phenome-wide association plot on 258 ordinal categorical phenotypes, (B) regional association plots between rs139315125, (C) sample size distribution in different categories, (D) minor allele frequencies in different categories

**Figure S15.**
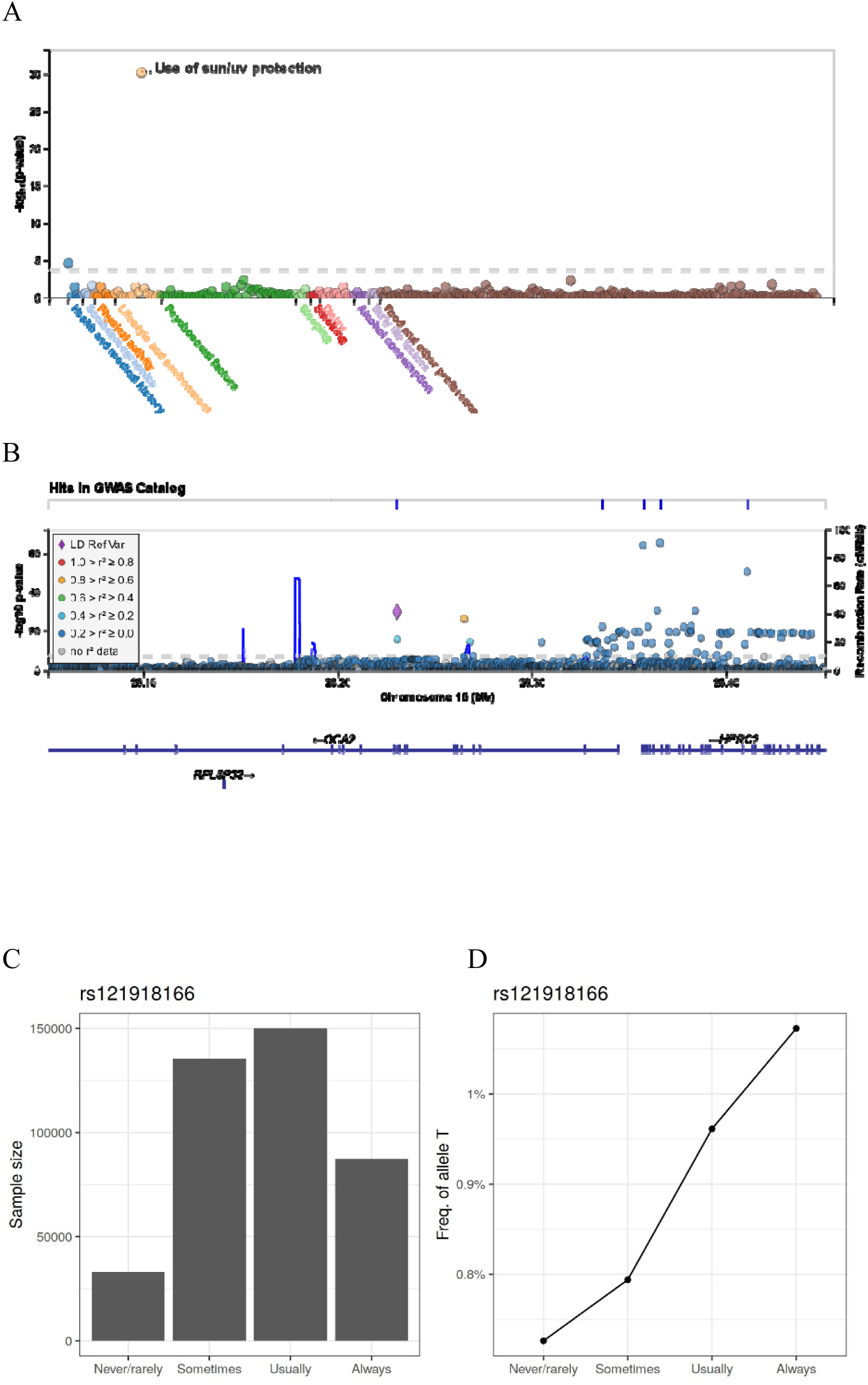
Use of sun/uv protection versus rs121918166, a nonsynonymous single nucleotide variant in OCA2 Gene. The phenotypes (Field ID: 2267) include 4 ordinal categories: Never/rarely, sometimes, most of the time, and always. (A) phenome-wide association plot on 258 ordinal categorical phenotypes, (B) regional association plots between rs121918166, (C) sample size distribution in different categories, (D) minor allele frequencies in different categories

**Figure S16.**
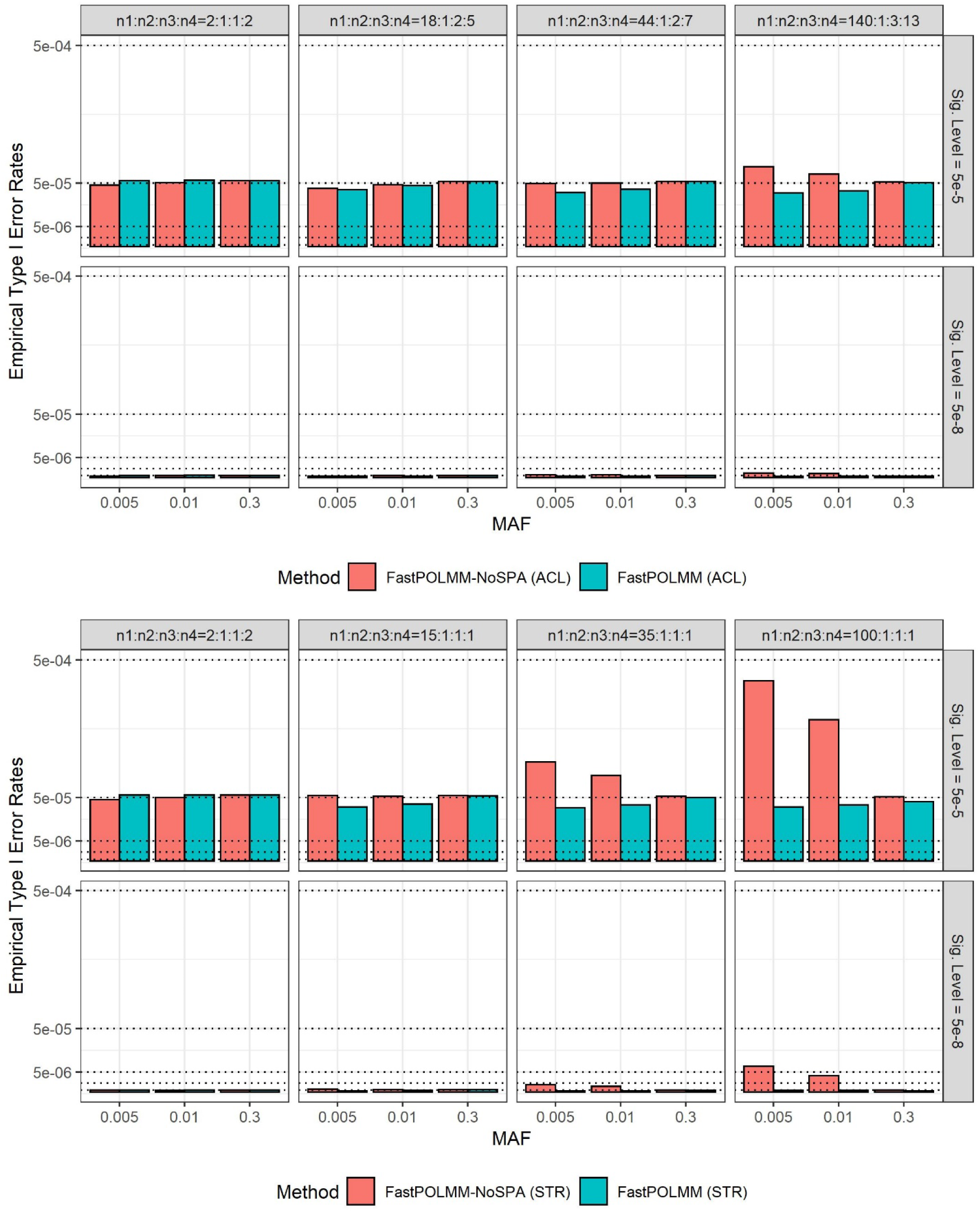
Type I error rates of FastPOLMM methods when categorical phenotypes were simulated following adjacent category logistic model (ACL, upper panels) and stereotype model (STR, lower panels). For both ACL and STR, we evaluated type I error rates at significance levels of 5e-5 and 5e-8. More details about the ACL and STR models and simulations can be seen in Supplementary Note.

**Table S1.** Comparison of different methods for computation time and memory usage given different sample sizes. CPU core is Intel(R) Xeon(R) Gold 6138 @ 2.00GHz.

| Sample Size | BOLT-LMM | SAIGE | DensePOLMM |  | FastPOLMM |  | fastGWA |
| --- | --- | --- | --- | --- | --- | --- | --- |
|  |  |  | 3 levels | 6 levels | 3 levels | 6 levels |  |
| A. Computation time in Step 1 (Hours) |  |  |  |  |  |  |  |
| 10000 | 0.1411 | 0.4177 | 0.3630 | 0.3712 | 0.0045 | 0.0049 | 0.0007 |
| 20000 | 0.2563 | 0.7367 | 0.6042 | 0.7134 | 0.0048 | 0.0102 | 0.0014 |
| 40000 | 0.4649 | 1.2081 | 1.2237 | 1.3909 | 0.0062 | 0.0162 | 0.0015 |
| 1.00E+05 | 1.4462 | 3.6556 | 5.9188 | 7.3769 | 0.0124 | 0.0241 | 0.0038 |
| 2.00E+05 | 3.4735 | 9.6960 | 18.1085 | 21.2003 | 0.0223 | 0.0406 | 0.0090 |
| 397798 | 8.7587 | 40.3603 | 49.9075 | 64.1850 | 0.0324 | 0.0893 | 0.0203 |
| B. Computation time in Step 2 (Hours) |  |  |  |  |  |  |  |
| 10000 | 58.845 | 83.098 | 61.616 | 61.476 | 61.616 | 61.476 | 39.229 |
| 20000 | 70.274 | 93.073 | 69.583 | 69.664 | 69.583 | 69.664 | 39.753 |
| 40000 | 79.088 | 110.238 | 89.650 | 89.668 | 89.650 | 89.668 | 42.461 |
| 1.00E+05 | 95.520 | 184.972 | 146.928 | 155.909 | 146.928 | 155.909 | 49.548 |
| 2.00E+05 | 109.971 | 310.370 | 240.802 | 261.855 | 240.802 | 261.855 | 64.996 |
| 397798 | 118.605 | 384.618 | 296.906 | 339.898 | 296.906 | 339.898 | 81.112 |
| C. Memory usage (GB) |  |  |  |  |  |  |  |
| 10000 | 1.684 | 1.483 | 1.378 | 1.465 | 1.168 | 1.208 | < 0.400 |
| 20000 | 2.480 | 2.349 | 2.002 | 2.057 | 1.167 | 1.250 | < 0.400 |
| 40000 | 4.098 | 4.072 | 3.284 | 3.417 | 1.228 | 1.316 | < 0.400 |
| 1.00E+05 | 9.028 | 8.912 | 7.060 | 7.333 | 1.332 | 1.863 | < 0.400 |
| 2.00E+05 | 17.378 | 17.199 | 13.453 | 14.128 | 1.774 | 2.511 | 0.580 |
| 397798 | 33.996 | 33.513 | 25.986 | 27.700 | 2.561 | 4.140 | 0.870 |

**Table S2.** Summary of the 258 ordinal categorical phenotypes in UK Biobank data analysis

| A. Distribution of different phenotype categories |  | B. Distribution of level numbers |  |
| --- | --- | --- | --- |
| Phenotype category | Number | Level number | Number |
| Food and other preference | 150 | 3 | 20 |
| Psychosocial factors | 46 | 4 | 34 |
| Lifestyle and environment | 16 | 5 | 23 |
| Dietry | 12 | 6 | 23 |
| Physical activity | 7 | 7 | 5 |
| Alcohol consumption | 5 | 8 | 3 |
| Health and medical history | 5 | 9 | 150 |
| Sleeping | 5 |  |  |
| Early life factors | 4 |  |  |
| Smoking | 4 |  |  |
| Sociodemographics | 4 |  |  |

**Table S3.** Summary of markers used in data analysis of “Liking for adding salt to foods”. Gene annotation and Polyphen2 HDIV score are from ANNOVAR. Based on Polyphen2 HDIV score, the nonsynonymous variants are divided into 3 groups: Probably damaging variants (Polyphen2 HDIV score ≥ 0.957), Possibly damaging variants (Polyphen2 HDIV score ≥ 0.453 and ≤ 0.956), and Benign variants (Polyphen2 HDIV score ≤ 0.452).

| Variants in genome | Exon variants | Nonsynonymous variants (proportion in exon variants) | Probably damaging variants (proportion in nonsynonymous variants) | Possibly damaging variants (proportion in nonsynonymous variants) | Benign variants (proportion in nonsynonymous variants) |
| --- | --- | --- | --- | --- | --- |
| All variants in data analysis |  |  |  |  |  |
| 24,135,906 | 229,586 | 124,975 (54.4%) | 41,092 (32.9%) | 20,141 (16.1%) | 58,450 (46.8%) |
| Significant variants for 258 phenotypes (p value < 5e-8) |  |  |  |  |  |
| 5,885 | 275 | 207 (75.3%) | 63 (30.4%) | 33 (15.9%) | 111 (53.6%) |

## Notes

### Competing Interest Statement

The authors have declared no competing interest.

