## Supplementary Note for "Efficient mixed model approach for large-scale genome-wide association studies of ordinal categorical phenotypes"

1. **Algorithm details**
   1.1. **Maximum Likelihood Estimation of POLMM.** For mathematical convenience, we define a $J\times1$ vector $\tilde{y}_{i}=\left( y_{i1},\cdots,y_{iJ} \right)^{T}$ as an equivalent representation of the ordinal categorical phenotype $y_{i}$ : if $y_{i}=j$, then $y_{ij}=1$ and the other elements in $\tilde{y}_{i}$ are 0. For subject $i$, the conditional log-likelihood function given random effects $b$ is

$$l_{i}\left( \beta,\gamma;b,\epsilon\right)=\log\left( \Pr\left( y_{i} \right) \right)=\sum_{j=1}^{J} y_{ij}\log\left( \mu_{ij} \right),$$

where $\mu_{ij}$ is the mean of $y_{ij}$, that is,

$$\mu_{ij}=E\left( y_{ij} \right)=\Pr(y_{ij}=1)=\Pr(y_{i}=j)=\Pr(y_{i}\leq j)-\Pr(y_{i}\leq j-1).$$

The first partial derivative of $l_{i}\left( \beta,\gamma;b,\epsilon\right)$ with respect to the linear predicator $\eta_{i}$ is

$$\frac{\partial l_{i}\left( \beta,\gamma;b,\epsilon\right)}{\partial\eta_{i}}=\sum_{j=1}^{J} \frac{y_{ij}}{\mu_{ij}}\cdot\frac{\partial\mu_{ij}}{\partial\eta_{i}}=\sum_{j=1}^{J} \frac{\left( y_{ij}-\mu_{ij} \right)}{\mu_{ij}}\cdot\frac{\partial\mu_{ij}}{\partial\eta_{i}}=\sum_{j=1}^{J-1} \frac{\left( y_{ij}-\mu_{ij} \right)}{\mu_{ij}}\cdot\frac{\partial\mu_{ij}}{\partial\eta_{i}}+\frac{\left( y_{iJ}-\mu_{iJ} \right)}{\mu_{iJ}}\cdot\frac{\partial\mu_{iJ}}{\partial\eta_{i}}=\sum_{j=1}^{J-1} \frac{\left( y_{ij}-\mu_{ij} \right)}{\mu_{ij}}\cdot\frac{\partial\mu_{ij}}{\partial\eta_{i}}-\sum_{j=1}^{J-1} \frac{\left( y_{ij}-\mu_{ij} \right)}{\mu_{iJ}}\cdot\frac{\partial\mu_{iJ}}{\partial\eta_{i}}=\sum_{j=1}^{J-1} \left( y_{ij}-\mu_{ij} \right)\cdot\left[ \frac{1}{\mu_{ij}}\cdot\frac{\partial\mu_{ij}}{\partial\eta_{i}}-\frac{1}{\mu_{iJ}}\cdot\frac{\partial\mu_{iJ}}{\partial\eta_{i}} \right].$$

The second and fourth equations hold since $\sum_{j=1}^{J} y_{ij}=\sum_{j=1}^{J} \mu_{ij}=1$ and$\sum_{j=1}^{J} \partial\mu_{ij}/\partial\eta_{i}=0.$ We define $n\left( J-1 \right)\times1$ vectors

$$\tilde{y}=\left( y_{11},y_{12},\cdots,y_{1\left( J-1 \right)},y_{21},y_{22},\cdots,y_{2\left( J-1 \right)},\cdots,y_{n1},y_{n2},\cdots,y_{n\left( J-1 \right)} \right)^{T},$$

$$\tilde{\mu}=\left( \mu_{11},\mu_{12},\cdots,\mu_{1\left( J-1 \right)},\mu_{21},\mu_{22},\cdots,\mu_{2(J-1)},\cdots,\mu_{n1},\mu_{n2},\cdots,\mu_{n\left( J-1 \right)} \right)^{T},$$

an $n\left( J-1 \right)\times n\left( J-1 \right)$ diagonal matrix

$$R=diag\left( R_{11},R_{12},\cdots,R_{1\left( J-1 \right)},R_{21},R_{22},\cdots,R_{2\left( J-1 \right)},\cdots,R_{n1},R_{n2},\cdots,R_{n\left( J-1 \right)} \right)$$

where$R_{ij}=1/\mu_{ij}\cdot\partial\mu_{ij}/\partial\eta_{i}-1/\mu_{iJ}\cdot\partial\mu_{iJ}/\partial\eta_{i}$, and an $n\left( J-1 \right)\times n$ matrix

$$\tilde{Z}=\left( e_{1},\cdots,e_{1},e_{2},\cdots,e_{2}\cdots,e_{n},\cdots,e_{n} \right)^{T}$$

where $e_{i}$ denotes an $n\times1$ vector with a 1 in the $i$-th coordinate and 0’s elsewhere. Then, the first derivative of log-likelihood function $l\left( \beta,\gamma;b,\epsilon\right)=\sum_{i\leq n} l_{i}\left( \beta,\gamma;b,\epsilon\right)$ with respect to $\eta=\left( \eta_{1},\cdots,\eta_{n} \right)^{T}$ is

$$\frac{\partial l\left( \beta,\gamma;b,\epsilon\right)}{\partial\eta}=\tilde{Z}^{T}R\left( \tilde{y}-\tilde{\mu} \right),$$

and the first derivatives of $l\left( \beta,\gamma;b,\epsilon\right)$ with respect to $\left( \beta,\gamma,b \right)$ are

$$\frac{\partial l\left( \beta,\gamma;b,\epsilon\right)}{\partial\beta}={X^{T}\tilde{Z}}^{T}R\left( \tilde{y}-\tilde{\mu} \right), \frac{\partial l\left( \beta,\gamma;b,\epsilon\right)}{\partial\gamma}=G^{T}\tilde{Z}^{T}R\left( \tilde{y}-\tilde{\mu} \right), \frac{\partial l\left( \beta,\gamma;b,\epsilon\right)}{\partial b}=\tilde{Z}^{T}R\left( \tilde{y}-\tilde{\mu} \right).$$

We let an $n\left( J-1 \right)\times n\left( J-1 \right)$ block diagonal matrix $\Psi$ denote the covariance matrix of $\tilde{y}$ as follows.

$$\Psi=\left[ \begin{matrix} \Psi_{1} & 0 & 0 \\ 0 & \ddots& 0 \\ 0 & 0 & \Psi_{n} \end{matrix} \right], \Psi_{i}=\left[ \begin{matrix} \mu_{i1} & 0 & 0 \\ 0 & \ddots& 0 \\ 0 & 0 & \mu_{i\left( J-1 \right)} \end{matrix} \right]-\mu_{i}\mu_{i}^{T}, \mu_{i}=\left( \mu_{i1},\cdots,\mu_{i\left( J-1 \right)} \right)^{T}.$$

Then, under certain regularity conditions^1^, the second derivative of $l\left( \beta,\gamma;b,\epsilon\right)$ with respect to $b$ can be approximated by

$$\frac{\partial^{2}l\left( \beta,\gamma;b,\epsilon\right)}{\partial b\partial b^{T}}\approx E\left( \frac{\partial^{2}l\left( \beta,\gamma;b,\epsilon\right)}{\partial b\partial b^{T}} \right)\approx E\left( -\frac{\partial l\left( \beta,\gamma;b,\epsilon\right)}{\partial b}\frac{\partial l\left( \beta,\gamma;b,\epsilon\right)}{\partial b^{T}} \right)=-\tilde{Z}^{T}R\cdot E\left( \left( \tilde{y}-\tilde{\mu} \right)\cdot\left( \tilde{y}-\tilde{\mu} \right)^{T} \right)\cdot R\tilde{Z}=-\tilde{Z}^{T}R\Psi R\tilde{Z}.$$

We assume that random vector $b$ follows a multivariate normal distribution $N(0,\tau V)$, then the marginal log-likelihood function of $(\beta,\gamma,\tau)$ is

$$l\left( \beta,\gamma,\tau;\epsilon\right)=\log\int\exp\left\{ l\left( \beta,\gamma;b,\epsilon\right) \right\}\times\left( 2\pi\right)^{-\frac{n}{2}}\left| \tau V \right|^{-\frac{1}{2}}\times\exp\left\{ -\frac{1}{2}b^{T}\left( \tau V \right)^{-1}b \right\}db.$$

Similar to GMMAT^2^, we use Laplace method to approximate the $n$-dimensional integral, and the marginal log-likelihood function becomes

$l\left( \beta,\gamma,\tau;\epsilon\right)\approx-\frac{1}{2}\log\left| \tau V \right|-\frac{1}{2}\log\left| -f^{''}\left( \tilde{b} \right) \right|+f(\tilde{b})$	(2)

where

$f\left( b \right)=l\left( \beta,\gamma;b,\epsilon\right)-\frac{1}{2}b^{T}\left( \tau V \right)^{-1}b$, $\tilde{b}=\arg\max f(b)$,

and the second derivative

$$f^{''}\left( b \right)=\frac{\partial^{2}l\left( \beta,\gamma;b,\epsilon\right)}{\partial b\partial b^{T}}-\left( \tau V \right)^{-1}\approx-\tilde{Z}^{T}R\Psi R\tilde{Z}-\left( \tau V \right)^{-1}.$$

Following GMMAT^2^ and SAIGE^3^, we assume that matrix $R$and$\Psi$ change slowly with respect to $\eta$. The derivatives of equation (2) with respect to $\left( \beta,\gamma,b \right)$ are

$$\frac{\partial l\left( \beta,\gamma,\tau;\epsilon\right)}{\partial\beta}=\frac{\partial l\left( \beta,\gamma;b,\epsilon\right)}{\partial\beta}={X^{T}\tilde{Z}}^{T}R\left( \tilde{y}-\tilde{\mu} \right),$$

$$\frac{\partial l\left( \beta,\gamma,\tau;\epsilon\right)}{\partial\gamma}=\frac{\partial l\left( \beta,\gamma;b,\epsilon\right)}{\partial\gamma}={G^{T}\tilde{Z}}^{T}R\left( \tilde{y}-\tilde{\mu} \right),$$

$$\frac{\partial l\left( \beta,\gamma,\tau;\epsilon\right)}{\partial b}=\frac{\partial l\left( \beta,\gamma;b,\epsilon\right)}{\partial b}-\left( \tau V \right)^{-1}b=\tilde{Z}^{T}R\left( \tilde{y}-\tilde{\mu} \right)-\left( \tau V \right)^{-1}b.$$

Under the null hypothesis $\gamma=0$, if $\epsilon$ and $\tau$ are known, we jointly choose $\hat{\beta}(\epsilon,\tau)$ and $\hat{b}(\epsilon,\tau)$ to maximize $l\left( \beta,\gamma,\tau;\epsilon\right)$, then $\hat{b}\left( \epsilon,\tau\right)=\tilde{b}(\hat{\beta}\left( \epsilon,\tau\right),\gamma=0)$ because $\tilde{b}$ maximizes $f\left( b \right)$ for given $(\beta,\gamma)$.^2^ Defining a working vector $\tilde{Y}=\tilde{Z}\eta+R^{-1}\Psi^{-1}\left( \tilde{y}-\tilde{\mu} \right)$, the solution of

$${X^{T}\tilde{Z}}^{T}R\left( \tilde{y}-\tilde{\mu} \right)=0, \tilde{Z}^{T}R\left( \tilde{y}-\tilde{\mu} \right)-\left( \tau V \right)^{-1}b=0,$$

can be written as the solution to the system

$$\left[ \begin{matrix} {X^{T}\tilde{Z}}^{T}R\Psi R\tilde{Z}X & {X^{T}\tilde{Z}}^{T}R\Psi R\tilde{Z} \\ \tilde{Z}^{T}R\Psi R\tilde{Z}X & \tilde{Z}^{T}R\Psi R\tilde{Z}+\left( \tau V \right)^{-1} \end{matrix} \right]\left[ \begin{matrix} \beta\\ b \end{matrix} \right]=\left[ \begin{matrix} {X^{T}\tilde{Z}}^{T}R\Psi R\tilde{Y} \\ \tilde{Z}^{T}R\Psi R\tilde{Y} \end{matrix} \right]$$

Let $\tilde{V}=\tilde{Z}V\tilde{Z}^{T}$, $\Sigma=R^{-1}\Psi^{-1}R^{-1}+\tau\tilde{V}$, and $P=\Sigma^{-1}-\Sigma^{-1}\tilde{Z}X\left( {X^{T}\tilde{Z}}^{T}\Sigma^{-1}\tilde{Z}X \right)^{-1}{X^{T}\tilde{Z}}^{T}\Sigma^{-1}$, then

$$\hat{\beta}=\left( {X^{T}\tilde{Z}}^{T}\Sigma^{-1}\tilde{Z}X \right)^{-1}{X^{T}\tilde{Z}}^{T}\Sigma^{-1}\tilde{Y}, \hat{b}=\tau V\cdot\tilde{Z}^{T}\Sigma^{-1}\left( \tilde{Y}-\tilde{Z}X\hat{\beta} \right) (2)$$

is the solution. We note that

$$\tilde{Y}-\tilde{Z}\eta=\tilde{Y}-\tilde{Z}X\hat{\beta}-\tilde{Z}\hat{b}=\left\{ I-\tau\tilde{V}\cdot\Sigma^{-1} \right\}\left( \tilde{Y}-\tilde{Z}X\hat{\beta} \right)=R^{-1}\Psi^{-1}R^{-1}\Sigma^{-1}\cdot\left( \tilde{Y}-\tilde{Z}X\hat{\beta} \right)=R^{-1}\Psi^{-1}R^{-1}P\tilde{Y}.$$

We add one intercept term with all elements of 1 to the covariate matrix and fix the first cutpoint $\epsilon_{1}=0$. Then, after updating $\hat{\beta}$ and $\hat{b}$, we use Newton–Raphson method to iteratively estimate cutpoints $\epsilon_{2},\cdots,\epsilon_{J-1}$ until convergence.

1.2. **Estimation of Variance Component.** Given random effect $\hat{b}$, vector $\tilde{y}$ has a mean of $\tilde{\mu}$ and a covariance matrix of $\Psi$. Using quasi-likelihood and Pearson chi-square statistics^4^, we approximate the log-likelihood

$$l\left( \beta,\gamma;\hat{b},\epsilon\right)\approx C_{1}-\frac{1}{2}\cdot\left( \tilde{y}-\tilde{\mu} \right)^{T}\Psi^{-1}\left( \tilde{y}-\tilde{\mu} \right)=C_{1}-\frac{1}{2}\cdot\left( \tilde{Y}-\tilde{Z}\eta\right)^{T}R\Psi R\left( \tilde{Y}-\tilde{Z}\eta\right),$$

where $C_{1}$ is independent from random vector $\tilde{y}$. Then, the log-likelihood function

$$l\left( \beta,\gamma,\tau;\epsilon\right)\approx-\frac{1}{2}\log\left| \tau V \right|-\frac{1}{2}\log\left| -f^{''}\left( \tilde{b} \right) \right|+f\left( \tilde{b} \right)\approx-\frac{1}{2}\log\left| \tau V \right|-\frac{1}{2}\log\left| \tilde{Z}^{T}R\Psi R\tilde{Z}+\left( \tau V \right)^{-1} \right|+l\left( \beta,\gamma;\hat{b},\epsilon\right)-\frac{1}{2}\hat{b}^{T}\left( \tau V \right)^{-1}\hat{b}\approx-\frac{1}{2}\log\left| I_{n}+\tau V\cdot\tilde{Z}^{T}R\Psi R\tilde{Z} \right|+C_{1}-\frac{1}{2}\cdot\left( \tilde{Y}-\tilde{Z}\eta\right)^{T}R\Psi R\left( \tilde{Y}-\tilde{Z}\eta\right)-\frac{1}{2}\left( \tilde{Y}-\tilde{X}\hat{\beta} \right)^{T}\Sigma^{-1}\tilde{Z}\cdot\left( \tau V \right)\cdot\tilde{Z}^{T}\Sigma^{-1}\left( \tilde{Y}-\tilde{X}\hat{\beta} \right)=-\frac{1}{2}\log\left| I_{n}+\tau\tilde{V}\cdot R\Psi R \right|+C_{1}-\frac{1}{2}\tilde{Y}^{T}PR^{-1}\Psi^{-1}R^{-1}P\tilde{Y}-\frac{1}{2}\tilde{Y}^{T}P\cdot\left( \tau\tilde{V} \right)\cdot P\tilde{Y}=-\frac{1}{2}\log\left| \left( R^{-1}\Psi^{-1}R^{-1}+\tau\tilde{V} \right)\cdot R\Psi R \right|+C_{1}-\frac{1}{2}\tilde{Y}^{T}P\left( R^{-1}\Psi^{-1}R^{-1}+\tau\tilde{V} \right)P\tilde{Y}=-\frac{1}{2}\log\left| \Sigma\cdot R\Psi R \right|+C_{1}-\frac{1}{2}\tilde{Y}^{T}P\Sigma P\tilde{Y}=-\frac{1}{2}\log\left| \Sigma\right|-\frac{1}{2}\log\left| R\Psi R \right|+C_{1}-\frac{1}{2}\tilde{Y}^{T}P\tilde{Y}=C-\frac{1}{2}\log\left| \Sigma\right|-\frac{1}{2}\tilde{Y}^{T}P\tilde{Y}.$$

The restricted maximum likelihood (REML) version^2^ is

$$l_{R}\left( \beta,\gamma,\tau;\epsilon\right)\approx C_{R}-\frac{1}{2}\log\left| \Sigma\right|-\frac{1}{2}\log\left| \tilde{X}^{T}\Sigma^{-1}\tilde{X} \right|-\frac{1}{2}\tilde{Y}^{T}P\tilde{Y}$$

Since $\partial P/\partial\tau=-P\tilde{V}P$, the derivative

$$\frac{\partial l_{R}\left( \beta,\gamma,\tau;\epsilon\right)}{\partial\tau}=\frac{1}{2}\tilde{Y}^{T}P\tilde{V}P\tilde{Y}-\frac{1}{2}\mathrm{tr} \left[ P\tilde{V} \right],$$

and the average information

$$AI=\frac{1}{2}\cdot\tilde{Y}^{T}P\tilde{V}P\tilde{V}P\tilde{Y}.$$

We use the following workflow to fit the null POLMM:

1. Fit a proportional odds logistic model with $\tau=0$ and $\gamma=0$ to estimate $\hat{\beta}^{\left( 0 \right)},\hat{\epsilon}^{\left( 0 \right)}$, and then calculate $\tilde{Y}^{\left( 0 \right)}$. Set initial value $\hat{\tau}^{\left( 0 \right)}=0.2$.
2. Update $\hat{\beta}^{\left( 1 \right)},\hat{b}^{\left( 1 \right)}$ and $\hat{\epsilon}^{\left( 1 \right)}$ using $\hat{\tau}^{\left( 0 \right)}$ and $\tilde{Y}^{\left( 0 \right)}$;
   2.1. Update $\hat{\beta},\hat{b}$ following equation (2);
   2.2. Use Newton-Raphson algorithm to update $\hat{\epsilon}$ until converges;
   2.3. Repeat steps 2.1-2.2 until $\hat{\beta}$ converges;
3. Update $\tilde{Y}^{\left( 1 \right)}$ and $\hat{\tau}^{\left( 1 \right)}=\hat{\tau}^{\left( 0 \right)}+\left\{ AI^{\left( 1 \right)} \right\}^{-1}\left( \partial l_{R}(\hat{\tau}^{\left( 0 \right)})/\partial\tau\right)$ using $\hat{\beta}^{\left( 1 \right)},\hat{b}^{\left( 1 \right)}$and $\hat{\epsilon}^{\left( 1 \right)}$;
4. Repeat steps 2-3 until $\hat{\tau}$ converges.
   1.3. **Score Test and Saddlepoint Approximation.** We calculate an $n$-dimensional covariate-adjusted genotype vector $\bar{G}=G-X\left( {X^{T}\tilde{Z}}^{T}R\Psi R\tilde{Z}X \right)^{-1}{X^{T}\tilde{Z}}^{T}R\Psi R\tilde{Z}G.$Under the null hypothesis, the score statistic

$$T=\frac{\partial l\left( \beta,\tau;\epsilon\right)}{\partial\gamma}=G^{T}\tilde{Z}^{T}R\left( \tilde{y}-\tilde{\mu} \right)=\bar{G}^{T}\tilde{Z}^{T}R\left( \tilde{y}-\tilde{\mu} \right)=\bar{G}^{T}\tilde{Z}^{T}R\Psi R\left( \tilde{Y}-\tilde{\eta} \right)=\bar{G}^{T}\tilde{Z}^{T}P\tilde{Y}.$$

Since $\tilde{Y}=\tilde{Z}\eta+R^{-1}\Psi^{-1}\left( \tilde{y}-\tilde{\mu} \right)$, its estimated variance is

$$\hat{Var}\left( T \right)=E\left( \bar{G}^{T}\tilde{Z}^{T}P\tilde{Y}\tilde{Y}^{T}P\tilde{Z}\bar{G} \right)=\bar{G}^{T}\tilde{Z}^{T}P\cdot\left[ R^{-1}\Psi^{-1}\cdot E\left( \left( \tilde{y}-\tilde{\mu} \right)\cdot\left( \tilde{y}-\tilde{\mu} \right)^{T} \right)\cdot\Psi^{-1}R^{-1}+\tilde{Z}\cdot E(\eta\cdot\eta^{T})\cdot\tilde{Z}^{T} \right]\cdot P\tilde{Z}\bar{G}=\bar{G}^{T}\tilde{Z}^{T}P\cdot\left[ R^{-1}\Psi^{-1}R^{-1}+\tilde{Z}\cdot(\tau V)\cdot\tilde{Z}^{T} \right]\cdot P\tilde{Z}\bar{G}=\bar{G}^{T}\tilde{Z}^{T}P\Sigma P\tilde{Z}\bar{G}=\bar{G}^{T}\tilde{Z}^{T}P\tilde{Z}\bar{G}.$$

To estimate $\hat{Var}(T)$, calculating $\Sigma^{-1}\bar{G}$ is required, which is computationally expensive for a genome-wide analysis. To reduce the computation cost, we use the same strategy as in BOLT-LMM^5^ and SAIGE^3^. Firstly, we use a small number of variants to calculate $\hat{Var}(T)$ and $\hat{Var}^{*}\left( T \right)=\bar{G}^{T}\tilde{Z}^{T}R\Psi R\tilde{Z}\bar{G}$, and estimate ratio $\hat{r}$ using the mean of $\hat{Var}(T)/\hat{Var}^{*}(T)$. Then, for each variant to test, we calculate $\hat{Var}^{*}\left( T \right)$ and then estimate $\hat{Var}\left( T \right)=\hat{r}\cdot\hat{Var}^{*}\left( T \right)$. When estimating $\hat{r}$, we increase the number of variants until the coefficient of variation for the ratio estimation is lower than a pre-given cutoff of 0.0025. In both simulation studies and real data analysis, the variant number is usually less than 30, which is consistent with SAIGE.

For each variant, the variance-adjusted test statistics is

$$T_{adj}=\frac{T}{\sqrt{\hat{Var}(T)}}=\frac{\bar{G}^{T}\tilde{Z}^{T}R\left( \tilde{y}-\tilde{\mu} \right)}{\sqrt{\hat{r}\bar{G}^{T}\tilde{Z}^{T}R\Psi R\tilde{Z}\bar{G}}},$$

which has mean zero and variance one under the null hypothesis. The regular score test assumes that $T_{adj}$ asymptotically follows a standard normal distribution, which uses only the first two moments. However, when the sample size distribution of different categories is highly unbalanced, the underlying distribution of $T_{adj}$ could be substantially different from a standard normal distribution, especially when testing low-frequency variants. To accurately calculate p values, we use saddlepoint approximation which uses the entire cumulant generating function (CGF) to approximates the null distribution. Suppose that $\bar{G}_{i}$ is the $i$-th element in vector $\bar{G}$, we define

$$T_{i}=\sum_{j=1}^{J-1} \frac{\bar{G}_{i}R_{ij}\left( y_{ij}-\mu_{ij} \right)}{\sqrt{\bar{G}^{T}R\Psi R\bar{G}}}=\sum_{j=1}^{J-1} c_{ij}y_{ij}-\sum_{j=1}^{J-1} c_{ij}\mu_{ij}, c_{ij}=\frac{\bar{G}_{i}R_{ij}}{\sqrt{\bar{G}^{T}R\Psi R\bar{G}}},$$

then the statistic

$$T_{adj}=\frac{\bar{G}^{T}R\left( \tilde{y}-\tilde{\mu} \right)}{\sqrt{\hat{r}\bar{G}^{T}R\Psi R\bar{G}}}=\frac{1}{\sqrt{\hat{r}}}\cdot\sum_{i=1}^{n} T_{i}.$$

Since $y_{ij}$ follows a Berounlli ($\mu_{ij}$) distribution and $\sum_{j} y_{ij}\leq1$, the CGF of $T_{i}$ is

$$K_{i}\left( t \right)=\log\left[ E\left( e^{tT_{i}} \right) \right]=\log\left( 1-\sum_{j=1}^{J-1} \mu_{ij}+\sum_{j=1}^{J-1} e^{c_{ij}t}\mu_{ij} \right)-\left( \sum_{j=1}^{J-1} c_{ij}\mu_{ij} \right)t,$$

and its derivatives

$$K_{i}^{'}\left( t \right)=\frac{\sum_{j=1}^{J-1} e^{c_{ij}t}\mu_{ij}c_{ij}}{1-\sum_{j=1}^{J-1} \mu_{ij}+\sum_{j=1}^{J-1} e^{c_{ij}t}\mu_{ij}}-\left( \sum_{j=1}^{J-1} c_{ij}\mu_{ij} \right),$$

$$K_{i}^{''}\left( t \right)=\frac{\left[ \sum_{j=1}^{J-1} e^{c_{ij}t}\mu_{ij}c_{ij}^{2} \right]\cdot\left[ 1-\sum_{j=1}^{J-1} \mu_{ij}+\sum_{j=1}^{J-1} e^{c_{ij}t}\mu_{ij} \right]-\left[ \sum_{j=1}^{J-1} e^{c_{ij}t}\mu_{ij}c_{ij} \right]^{2}}{\left[ 1-\sum_{j=1}^{J-1} \mu_{ij}+\sum_{j=1}^{J-1} e^{c_{ij}t}\mu_{ij} \right]^{2}}.$$

We use $K\left( t \right)=\sum_{i=1}^{n} K_{i}\left( t \right)$ to approximate CGF of $T_{adj}$ such that the variance from CGF is 1, that is,

$$K^{''}\left( 0 \right)=\sum_{i=1}^{n} K_{i}^{''}\left( 0 \right)=\sum_{i=1}^{n} \left\{ \sum_{j=1}^{J-1} \mu_{ij}c_{ij}^{2}-\left[ \sum_{j=1}^{J-1} \mu_{ij}c_{ij} \right]^{2} \right\}=\tilde{c}^{T}\Psi\tilde{c}=\frac{\bar{G}^{T}R\Psi R\bar{G}}{\bar{G}^{T}R\Psi R\bar{G}}=1,$$

where

$$\tilde{c}=\left( c_{11},c_{12},\cdots,c_{1\left( J-1 \right)},c_{21},c_{22},\cdots,c_{2\left( J-1 \right)},\cdots,c_{n1},c_{n2},\cdots,c_{n\left( J-1 \right)} \right)^{T}$$

The distribution of $T_{adj}$ at the observed test statistic $q$ can be approximated by

$$\Pr\left( T_{adj}<q \right)\approx F\left( q \right)=\Phi\left( w+\frac{1}{w}\log\left( \frac{v}{w} \right) \right),$$

where

$$w=sign\left( \hat{\zeta} \right)\sqrt{2\left\{ \hat{\zeta}q-K(\hat{\zeta}) \right\}}, v=\hat{\zeta}\sqrt{K''(\hat{\zeta})},$$

and $\hat{\zeta}$ is the solution of the equation $K^{'}\left( \hat{\zeta} \right)=q$.

We apply a hybrid strategy: if $\left| T_{adj} \right|<2$, p values are calculated based on normal approximation in which the variance is $\hat{Var}\left( T \right)=\hat{r}\cdot\hat{Var}^{*}\left( T \right)$; if $\left| T_{adj} \right|\geq2$, p values are calculated based on saddlepoint approximation. Using this hybrid strategy, we can greatly reduce computation time while controlling type I error rates. After fitting the null model, we calculate and store the following matrix

$$A_{1}=X\left( {X^{T}\tilde{Z}}^{T}R\Psi R\tilde{Z}X \right)^{-1}, A_{2}={X^{T}\tilde{Z}}^{T}R\Psi R\tilde{Z}, A_{3}=\tilde{Z}^{T}R\left( \tilde{y}-\tilde{\mu} \right), A_{4}=\tilde{Z}^{T}R\Psi R\tilde{Z}.$$

For each variant, it takes $O\left( np \right)$ computations to calculate vector $\bar{G}=G-A_{1}\cdot A_{2}\cdot G$. Because $A_{4}$ is a diagonal matrix, it takes $O\left( n \right)$ to calculate the score statistic $T=\bar{G}^{T}\cdot A_{3}$ and the variance $\hat{Var}^{*}\left( T \right)=\bar{G}^{T}A_{4}\bar{G}$. Thus, for normal distribution approximation, the computational complexity is still $O\left( np \right)$ and does not increase as the category number $J$ increases. For saddlepoint approximation, we use a partially normal approximation method to speed up the computation.^6^ Suppose that the first $m$ subjects have at least one minor allele each and the rest have homozygous major genotypes. We can express

$$T_{adj}=\frac{1}{\sqrt{\hat{r}}}\cdot\sum_{i=1}^{n} T_{i}=\frac{1}{\sqrt{\hat{r}}}\cdot\left( T_{\left( 1 \right)}+T_{\left( 2 \right)} \right)$$

where $T_{\left( 1 \right)}=\sum_{i=1}^{m} T_{i}$ and $T_{\left( 2 \right)}=\sum_{i=m+1}^{n} T_{i}$. Let $W=\left( {X^{T}\tilde{Z}}^{T}R\Psi R\tilde{Z}X \right)^{-1}{X^{T}\tilde{Z}}^{T}R\Psi R\tilde{Z}G$, and let $W_{l}$ be the $l$^th^ element of $W$. Then, we can further express $T_{\left( 2 \right)}$ as

$$T_{\left( 2 \right)}=\frac{1}{\sqrt{\hat{Var}^{*}\left( T \right)}}\cdot\sum_{i=m+1}^{n} \bar{G}_{i}\left( \sum_{j=1}^{J-1} R_{ij}\left( y_{ij}-\mu_{ij} \right) \right)=\frac{1}{\sqrt{\hat{Var}^{*}\left( T \right)}}\cdot\sum_{i=m+1}^{n} \left( 0-X_{i}W \right)\left( \sum_{j=1}^{J-1} R_{ij}\left( y_{ij}-\mu_{ij} \right) \right)=-\frac{1}{\sqrt{\hat{Var}^{*}\left( T \right)}}\cdot\sum_{i=m+1}^{n} \sum_{l=1}^{p} X_{il}W_{l}\left( \sum_{j=1}^{J-1} R_{ij}\left( y_{ij}-\mu_{ij} \right) \right)=-\frac{1}{\sqrt{\hat{Var}^{*}\left( T \right)}}\cdot\sum_{l=1}^{p} W_{l}\cdot\sum_{i=m+1}^{n} X_{il}\left( \sum_{j=1}^{J-1} R_{ij}\left( y_{ij}-\mu_{ij} \right) \right)=-\frac{1}{\sqrt{\hat{Var}^{*}\left( T \right)}}\cdot\sum_{l=1}^{p} W_{l}\cdot T_{\left( 2l \right)}$$

where

$$T_{\left( 2l \right)}=\sum_{i=m+1}^{n} \sum_{j=1}^{J-1} \left\{ X_{il}R_{ij}\left( y_{ij}-\mu_{ij} \right) \right\}.$$

If we assume that the non-genetic covariates are relatively balanced in the sample, then the normal approximation should be a good approximation of the null distribution of each $T_{\left( 2l \right)}$. Because $T_{\left( 2 \right)}$ is a weighted sum of the $T_{\left( 2l \right)}$ variables, we can also approximate the null distribution of $T_{\left( 2 \right)}$ by using a normal distribution and the CGF of $T_{\left( 2 \right)}$ can be approximated by

$$K_{\left( 2 \right)}(t)=\frac{1}{2}t^{2}\cdot V_{H_{0}}\left( T_{\left( 2 \right)} \right)$$

where

$$V_{H_{0}}\left( T_{\left( 2 \right)} \right)=\sum_{i=m+1}^{n} \frac{\bar{G}_{i}^{2}\cdot R_{i}^{T}\Psi_{i}R_{i}}{\hat{Var}^{*}\left( T \right)}$$

and $R_{i}=\left( R_{i1},R_{i2},\cdots,R_{i\left( J-1 \right)} \right)^{T}$. Hence, using the partially normal approximation, the CGF of $T_{adj}$ is $K\left( t \right)=\sum_{i=1}^{m} K_{i}\left( t \right)+K_{\left( 2 \right)}(t)$, and the saddlepoint approximation takes $O(m(J-1))$ computations to calculate the CGF and its derivatives.

2. **Data simulation of alternative models**

To evaluate the robustness of POLMM approaches, we also simulated categorical phenotypes following adjacent category logistic model (ACL) and stereotype model (STR). For subject $i\leq n$, the categorical phenotype $y_{i}=1,2,\ldots,J$ was simulated following

$$\log\left( \varsigma_{ij} \right)=\epsilon_{j}+\eta_{i}=\epsilon_{j}+X_{i}^{T}\beta+G_{i}\gamma+b_{i}, 1\leq j\leq J-1 (\mathrm{ACL})$$

where $\varsigma_{ij}=\Pr(y_{i}=j|X_{i},G_{i},b_{i})/\Pr(y_{i}=j+1|X_{i},G_{i},b_{i})$, and

$$\log\left( \varrho_{ij} \right)=\epsilon_{j}-\phi_{j}\eta_{i}=\epsilon_{j}-\phi_{j}\cdot\left( X_{i}^{T}\beta+G_{i}\gamma+b_{i} \right), 2\leq j\leq J (STR)$$

where $\varrho_{ij}=\Pr(y_{i}=j|X_{i},G_{i},b_{i})/\Pr(y_{i}=1|X_{i},G_{i},b_{i})$ and $\phi_{j}=j-1$. For both ACL and STR models, we simulated 10,000 subjects in 1,000 families based on the pedigree shown in Figure S3, in which each family included 10 subjects. We simulated covariates $X_{i}=\left( X_{i1},X_{i2} \right)^{T}$following a Bernoulli (0.5) distribution and the standard normal distribution. We simulated random effects $b=(b_{1},b_{2},\cdots,b_{n})$ following a multivariate normal distribution $N(0,\tau V)$ where variance component $\tau=1$ and $V$ is the real GRM from the family structure. We selected intercepts $\epsilon_{j}$ to simulate categorical phenotypes with different sample size distributions. Figure S15 shows the type I error rates results of POLMM approaches with and without using SPA under the null model $\gamma=0$ in different sample size distributions. For both ACL and STR models, POLMM method can still control type I error rates under significance levels of 5$\times{10}^{-5}$ and 5$\times{10}^{-8}$.
